# A New Feature of the Laboratory Model Plant *Nicotiana benthamiana*: Dead-End Trap for Sustainable Field Pest Control

**DOI:** 10.1101/2023.08.23.554404

**Authors:** Wen-Hao Han, Jun-Xia Wang, Feng-Bin Zhang, Shun-Xia Ji, Yu-Wei Zhong, Yin-Quan Liu, Shu-Sheng Liu, Xiao-Wei Wang

## Abstract

- Hemiptera and Thysanoptera insects pose persistent threats to agricultural production. Conventional management strategies involve the release of chemical or plastic agents, causing adverse environmental and global health issues. Notably, *Nicotiana benthamiana,* a globally utilized model plant, exhibits remarkable lethal effects and attraction towards these pests.
- In this study, we explored the potential of using *N. benthamiana* for Hemiptera and Thysanoptera pest control in the laboratory and field. Through net cover and three field assays over two years, we demonstrated the efficacy and benefits of using *N. benthamiana* as a field-deployed pest control dead-end trap.
- *N. benthamiana* demonstrated nearly 100% lethality to whiteflies, aphids, and thrips, with emitted volatiles attracting these insects. Field trials showed that potted and planted *N. benthamiana* blocks and traps whiteflies and thrips from several Solanaceae and Cucurbitaceae crops effectively, comparable to common commercial yellow and blue sticky boards. Moreover, *N. benthamiana* in the field exhibits robust growth in commercial greenhouses without negatively impacting crop growth, natural enemies, and pollinators.
- Our study introduces an innovative, easily implementable, and sustainable approach for controlling Hemiptera and Thysanoptera pests. Moreover, it unveils the novel utility of *N. benthamiana* in field-based pest management.

## Introduction

Conventional agricultural practices heavily rely on the use of chemical insecticides to combat herbivorous pests. However, these methods have severe and long-lasting negative consequences for ecosystems, global public health, and overall well-being (Tsiafouli *et al*., 2015; Tang *et al*., 2021). Compounding the issue, the emergence and evolution of insecticide resistance render pest control increasingly challenging, necessitating higher doses of pesticides to maintain yield (Kogan & Jepson, 2007; Sharma *et al*., 2019). Urgent action is needed to reduce insecticide usage, assuaging concerns about pesticide residues on food and the environmental pollution they cause (Candel *et al*., 2023).

In addition to insecticides, alternative methods such as sex pheromones, biotic agents, traps, and dead-trap plants play vital roles in pest control and global insecticide reduction, particularly in developing countries. The application of trap plants has emerged as an environmentally friendly alternative or complement to insecticides for managing insect pests. Traditional trap plants are more attractive to pests than the main crop, encouraging feeding and/or oviposition (Hokkanen, 1991; Godfrey & Leigh, 1994). Dead-end trap plants, on the other hand, possess heightened attractiveness to pests while being inhospitable to their survival or reproduction (Jackai & Singh, 1983; Thompson, 1988; Van Den Berg *et al*., 2001; Shelton & Nault, 2004; Badenes-Perez *et al*., 2005b; Shelton & Badenes-Perez, 2006; Khan *et al*., 2006). Incorporating dead-end trap plants into Integrated Pest Management (IPM) strategies offer promise in replacing commercial insecticides and sticky traps, thereby reducing the release of chemical and plastic pollutants into the environment (Shelton & Nault, 2004; Shelton & Badenes-Perez, 2006; Cook *et al*., 2007a). However, the development of novel dead-end trap plants is challenging, as the candidates must exhibit higher attractiveness and lethality to target pests than the main crop. Furthermore, some candidate plant species may not struggle to thrive in field environments, rendering them unsuitable for effective pest control. Consequently, the availability of field-appropriate and cost-effective dead-end trap plants remains limited. To date, several field-proven dead-end trap plants have demonstrated efficacy in controlling Lepidopteran, Coleoptera, and Diptera pests (Shelton & Badenes-Perez, 2006; Veromann *et al*., 2014; Ulmer *et al*., 2020; Gyawali *et al*., 2021; Rajesh *et al*., 2021; Yang *et al*., 2023). However, very few such plants have been reported for the management of other significant pests, such as Hemiptera and Thysanoptera insects.

Hemiptera and Thysanoptera insects pose continuous threats to agricultural yields, especially in vegetable crops. They inflict damage by sucking plant sap, leading to premature wilting, defoliation, stunted growth, and subsequent yield loss (Morse & Hoddle, 2006; Goggin, 2007; Walling, 2008; Wu *et al*., 2018). Additionally, many of these insects, including whiteflies, aphids, leafhoppers, and thrips, act as vectors for plant viruses and other pathogens causing yield losses and increasing the use of bactericides and fungicides in the field (Kaloshian & Walling, 2005; Gilbertson *et al*., 2015; Eigenbrode *et al*., 2018; Wang & Blanc, 2021). The management of Hemiptera and Thysanoptera insect pests has long relied on the use of pesticides, and the Arthropod Pesticide Resistance Database (http://www.pesticideresistance.org) has documented 3942 cases of insecticide resistance in Hemiptera and Thysanoptera insect pests (Guedes *et al*., 2016). However, to the best of our knowledge, no field-proven dead-end trap plants have been reported targeting Hemiptera and Thysanoptera insect pests.

*Nicotiana benthamiana* is widely utilized as a model plant in the study of plant innate immunity and defense signaling due to its suitability for virus-induced gene silencing and transient gene expression through leaf agroinfiltration (Goodin *et al*., 2008; Bally *et al*., 2015, 2018; Abrahamian *et al*., 2020; Rössner *et al*., 2022). Furthermore, in recent years, it has been employed as a plant bio-factory for the production of vaccines and pharmacological compounds (Arntzen, 2008, 2015; Larrimore *et al*., 2013; Lau & Sattely, 2015; Lomonossoff & D’Aoust, 2016; Martí *et al*., 2020). Notably, the *N. benthamiana* used in laboratories is native to the Granites site in the Northern Territory of Australia, an extremely remote and hostile location, exhibiting robust environmental resilience (Goodspeed, 1947; Bally *et al*., 2018). Numerous global studies employ *N. benthamiana* systems and its intricate genomic information is constantly being refined (Ranawaka *et al*., 2023). However, limited attention has been given to the interaction between *N. benthamiana* and insects. The responses and defense mechanisms of *N. benthamiana* against herbivorous insects remain unexplored. Moreover, the practical utility of *N.benthamiana* beyond laboratory settings, such as its potential as a dead-end trap, has yet to be investigated.

In this study, we present unexpected findings regarding the resistance and attractiveness of *N. benthamiana* compared to Solanaceae and Cucurbitaceae crops towards several Hemiptera and Thysanoptera insect pests. Furthermore, the laboratory model *N. benthamiana* plants were introduced into the field to control these pests. Greenhouse trials conducted over two years demonstrated that the potted and planted *N. benthamiana* plants serve as effective dead-end traps and viable alternatives to commercial sticky traps, facilitating pest control in the field pests. Moreover, *N. benthamiana* exhibits robust growth in commercial greenhouses without negatively impacting natural enemies.

## Materials and Methods

### Plant materials, insects, and experimental conditions

*Nicotiana benthamiana* LAB (laboratory isolate) strain, cultivated tobacco (*Nicotiana tabacum* cv. NC89), tomato (*Solanum lycopersicum* cv. Hezuo903), squash (*Cucurbita moschata* cv. Mibennangua), cucumber (*Cucumis sativus* cv. Jinyou616), eggplant (*Solanum melongena* cv. Zirong2), and capsicum (*Capsicum annuum* cv. Hangjiao1) were used in this study.

The whitefly cryptic species of *Bemisia tabaci* was MEAM1 (*mtCOI* GenBank: GQ332577). Whitefly adults within 7 days post-emergence were selected for the experiments. The green peach aphid *Myzus persicae* was reared on cultivated tobacco (*Nicotiana tabacum* cv. NC89). Winged adult aphids within 7 days post-emergence were selected for the experiments. The western flower thrips (*Frankliniella occidentalis*) and flower thrips (*Frankliniella intonsa*) were reared on cotton plants. The *Nesidiocoris tenuis* insects were reared on sesame plants. The experimental honeybees (*Apis mellifera*) were provided by the College of Animal Sciences at Zhejiang University. The experimental and insect-rearing plants were grown in a greenhouse with artificial illumination (14 h light:10 h dark cycle) and controlled the temperature (26 ± 2°C) and the relative humidity (60 ± 10%).

### Comparison of insect survival on different plants

The specially-designed nylon leaf cages (30 cm × 30 cm × 30 cm; 120 mesh) were used to evaluate the lethal effect on whiteflies, aphids, thrips, and *N. tenuis*. The leaf cages were fixed on the tested *N. benthamiana* plants and various other crops, and the tested pests (40 whiteflies, 40 aphids, 40 thrips, and 20 *N. tenuis*) were introduced into the cage individually. After 24 hours, the mortality of the pest was recorded. To evaluate the lethal effect on honeybees, an *N. benthamiana* plant was enclosed in a nylon cage with ten honeybees, and the survival rate was calculated after 48 hours. As a control, honeybees were placed in an empty cage.

### Choice bioassay

For the direct choice test, the comparison plants were positioned at both ends of a tailor-made nylon cage (length: 48 cm, width: 38 cm, height, 45 cm), and a glass petri dish (6 cm diameter) containing tested insect (100 whiteflies, 60 aphids, or 40 thrips) starved for 12 hours prior was placed equidistant from the two plants. After 6 hours, the number of insects settled on each plant was counted.

For the olfactometer test, the plants used for comparison were placed in two specially-made glass containers (Cylinder; 15.5 cm diameter × 51 cm height). Two streams of purified air, treated with activated charcoal, were directed through two glass containers into the olfactory arms at a flow of 3 L min^-1^. The olfactometer base was connected to a vacuum tube maintaining 6 L min^-1^ stable air-flow. Individual adult whitefly was released into the straight part at the base for the Y-tube (14.5 cm length of the base tube, 11.1 cm length of branch tube, 1cm diameter). Each insect was observed for up to 5 minutes, and a choice between two odor sources was recorded when the insect reached the end of either arm and remained there for at least 30 seconds. Each two odor sources were chosen by the tested insects, and these positions were interchanged after testing half of the insects to avert any interference of unanticipated asymmetries in the experimental setup.

### Net cover trials

The net cover trials were executed at Zhejiang University, Zijingang Campus, Hangzhou, China. The experimental site encompassed 12 rectangular plots (length: 4.09 m, width: 1.78 m). These plots had no chemical input for the past decade. The trials were conducted from July to October 2021.

The plots were leveled and covered with black waterproof plastic sheets. On top of these sheets, 200-mesh insect-proof nylon cages (length: 3.6 m, width: 2 m, height: 2 m) were installed. To ensure a controlled experimental environment, the surroundings were scrupulously purged of weeds or insects. The design of trial 1 followed the methodology outlined in the previous study (Cook *et al*., 2007b). In summary, the focal subjects were positioned upwind, while four designated points for insect release were strategically situated downwind. To simulate encounters during migration, a row of *N. benthamiana* plots (12 plots) was situated between these two distinct locations. Control for this plot involved the absence of *N. benthamiana* between the release point and the tomato plants **(Fig. S1a, b)**. Each release point was populated with 300 whiteflies, and the number of whiteflies on the tomato plants was conducted 24 hours post-experiment, with each plot comprising three replicates.

For trial 2, the same mesh infrastructure previously described was employed. Tomato plants were cultivated within nutrient bowls (length: 7 cm, width: 7 cm, height: 10 cm) for approximately 28 days, corresponding to the 4–5 leaf stage. A square arrangement of four potted *N. benthamiana* plants encircled the central tomato plants. A parallel control plot involved cultivating tomato plants devoid of *N. benthamiana*. Within each plot, four culture dishes were positioned at the four corners encircling the tomato plants, designated as release points for the whiteflies. These dishes were populated with 100 whiteflies each **(Fig. S1c, d)**. The number of whiteflies on the tomato plants was conducted 24 hours after release, serving as a metric for evaluating the mortality effect. Each setup had three replicates.

### Greenhouse trial 1 using N. benthamiana as a dead-end trap for pest control

Greenhouse trial 1 was executed at the same location and within the same period as the net cover trials. The experimental site was partitioned into 12 plots, and control and treatment plots were set up respectively, with three replicates. For the experimental period, indigenous whitefly populations naturally inhabited the experimental period, and these whiteflies were subsequently introduced from the environment to the targeted plants.

The trial layout adhered to the border design methodology previously documented (Javaid & Joshi, 1995). The focal tomato crops were encircled by 42 field-grown *N. benthamiana* plants. The cultivated *N. benthamiana* plants selected for the experiment were transplanted into the plots after approximately 21 days of growth within a controlled artificial climate chamber (3–4 leaves). The tomato seedlings were cultivated within similar conditions for about 14 days (3–4 leaves), and subsequently transplanted into the experimental plots surrounded by the cultivated *N. benthamiana* plants. A bi-daily enumeration of whitefly populations was conducted over a span of 19 days, alongside the observation and documentation of deceased whiteflies upon *N. benthamiana* leaves. During each counting session, all whiteflies present on every tomato plant within each plot were collected within a sampling container, subsequently undergoing a thorough enumeration. Control plots lacked *N. benthamiana* plants inclusion. **Fig. S2a, b** depict the schematic representation of the plot configurations, while **Fig. S2c** provides a comprehensive overview of the plot arrangement.

To elucidate the effect of potted *N. benthamiana* plants, tomato seedlings were employed as the experimental subject. Tomato seeds were sown within a 50-well tray (length: 25 cm, width: 50 cm, height: 5 cm). After 10 days, germinated tomato seedlings (2–3 leaves) were selected for the trial. Within each plot, a grouping of five trays, each harboring 50 tomato seedlings, was positioned at the central axis, enveloped by the presence of 42 potted *N. benthamiana* plants **(Fig. S2d, e)**. The schematic arrangement of the experimental setup can be found in **Fig. S2f**. The number of whiteflies on the tomato seedlings by the above way was counted every two days for the 13^th^ day.

### Greenhouse trial 2 using N. benthamiana as a substitute for commercial sticky traps

The Greenhouse trial 2 was conducted at the Zhejiang Academy of Agricultural Sciences. No fungicides or insecticides were applied within the experimental greenhouse. The experiment spanned from July to September 2022. Several greenhouse cubicles measuring 36 m^2 w^ere used in the trial. Each cubicle was divided into two plots by a grid (120 mesh) and two divided cubicles containing four plots were used for one repeat. Each plot contains six main crop plants and they were grown for 30 days, which naturally attracted whiteflies and thrips, but no other insects. Four plots were arranged with potted *N. benthamiana* plants along with yellow or blue sticky traps (30 cm × 25 cm) respectively, while the control plot remained without any additional settings. The positioning of the sticky traps and the *N. benthamiana* pots was set between two rows of main crops, 40–50 cm away from the plants on both sides **(Fig. S3)**. The height of the sticky traps was adjusted to align with the midpoint position of the *N. benthamiana* pots.

Observations were conducted at three-day intervals, during which the dead whiteflies or thrips adhered to the sticky traps or *N. benthamiana* pots. The number of whiteflies and thrips on the main crop leaves was recorded. The total number of whiteflies and thrips trapped by the *N. benthamiana* pots and commercial sticky traps in 15 days was counted and compared. The chosen 15-day duration represents the maximal effective operational period of the period during which the commercial sticky traps are employed in this study. After that, the surfaces of the sticky traps became susceptible to dust accumulation and water droplet adhesion, necessitating replacement. This process was repeated five times, thereby yielding five independent replicates. Tomatoes (*S*. *lycopersicum* cv. Hezuo903), peppers (*C*. *annuum* cv. Hangjiao1), or cucumbers (*C*. *sativus* cv. Jinyou616) served as the main crop in the above trials successively.

### Commercial greenhouse trials

The commercial greenhouse trials were conducted at the Yangdu base of the Zhejiang Academy of Agricultural Sciences in Jiaxing, Zhejiang, China. The experiment spanned from July to October 2022. The experimental greenhouse spanned an area of 160 m^2 (^20 m × 8 m) and was divided into five plots using a nylon net (120 mesh). Each plot consisted of four ridges covered with black plastic sheeting. Tomato plants were grown as the main crop and transplanted into the ridges at the stage of 3–4 leaves. Each tomato plant was placed 50 cm apart, and planted in two rows per ridge, resulting in a total of 48 tomato plants per plot. The plot settings were as follows: (1) potted *N. benthamiana* plants, (2) cultivated *N. benthamiana* plants, (3) commercial yellow sticky traps, (4) blue sticky traps, and (5) no additional setting (control). For setting 1, *N. benthamiana* plants were grown in pots (bottom diameter 12 cm, height 10 cm) and maintained in artificial climate chambers. They were then placed in the experimental plots at the stage of 8–9 leaves. A total of 16 potted *N. benthamiana* plants were evenly distributed in each experimental plot. For setting 2, the experimental *N. benthamiana* plants, cultivated in artificial climate chambers, were transplanted to the plots after approximately 21 days of growth (3–4 leaves). These transplanted *N. benthamiana* plants grew alongside the main crop, tomatoes. Again a total of 16 *N. benthamiana* plants were evenly distributed in each experimental plot. For setting 3 and 4, 16 commercial yellow or blue sticky traps (30 cm × 25 cm) were evenly placed within the plot, with a hanging height of 50–60 cm. In this experimental environment, the sticky traps needed to be replaced every 10 days to maintain their effectiveness. The replaced sticky traps were collected to prevent environmental pollution. The commercial greenhouse trials were conducted from August to October 2022, field statistics were recorded every seven days. The number of whiteflies and thrips on the main crop tomatoes as well as growth indicators, such as branch numbers, plant height, flowering, and fruiting, were counted. During this period, thrips were not observed in the field, resulting in the absence of data. Count one plant at every interval and a total of 24 plants were counted in each experimental plot. The setting for the commercial greenhouse trials is shown in **Fig. S4**.

### Statistical analysis

The differences in insect choice between the *N. benthamiana* plant and other crops were analyzed using nonparametric Wilcoxon matched pairs tests (for two dependent samples). In olfactometer tests, the differences in preference of tested insects and mites for the odor of the *N. benthamiana* plant and other crops were analyzed using the Binomial distribution test. The differences in mortality or survival rates of tested insects or mites on the *N. benthamiana* plant and other crops were analyzed with One-way ANOVA followed by Fisher’s least significant difference test using SPSS 26.0. Percentage data were arcsine square root transformed before analysis.

In net cover trial 1, the differences in whitefly numbers on the main tomato plants were analyzed using One-way ANOVA followed by Fisher’s least significant difference test. In net cover trial 2, the differences in whitefly numbers on the main tomato plants were analyzed using Student’s *t*-tests. In greenhouse trial 1, the differences in whitefly numbers on the main tomato plants were analyzed using Student’s *t*-tests. In greenhouse trial 2, the differences in the number of whiteflies and thrips trapped by the *N. benthamiana* plant and commercial sticky traps, as well as the number of whiteflies and thrips on the main crop tomatoes were analyzed using One-way ANOVA followed by Fisher’s least significant difference test. In the commercial greenhouse trials, the number of pests, natural enemies, and the trial outcomes of the main crop tomatoes were analyzed using One-way ANOVA followed by the Games-Howell test. SPSS 26.0 was used for the above analysis.

#### Image capture

Photos of the open field trials and commercial greenhouse trials were taken with iPhone 11 Pro Max (MWF42CH/A, Apple Inc., Cupertino, CA, USA). Dead whiteflies on the leaves of the *N. benthamiana* plant were photographed by a stereomicroscope imaging system (VHX-2000; KEYENCE AG, Osaka, Japan).

## Results

### *N. benthamiana* exhibits lethal effects on various Hemiptera and Thysanoptera insect pests

During our routine work in the greenhouse, we observed that the *N. benthamiana* plants often harbored numerous dead whiteflies **(boldFig. 1a)**, while other cultivated tobacco and vegetable crops in the vicinity had relatively few dead whiteflies. The dead whiteflies exhibited a dense distribution across the leaf surface, displaying evident distortion and desiccation **(Fig. 1b)**. Stereomicroscope examination revealed that the trichomes on the surface of *N. benthamiana* acted as a barrier, preventing whiteflies from accessing the leaf epidermis. Most of the whiteflies perished upon encountering the trichomes, being deprived of the opportunity to engage in feeding activities on the leaves **(Fig. 1c, d)**. To explore this phenomenon, we compared whitefly survival on *N. benthamiana* with six other crop species, including cultivated tobacco, tomato, capsicum, eggplant, squash, and cucumber. The mortality of whiteflies on *N. benthamiana* was nearly 100%, while it ranged mostly below 20% on the other tested plants **(Fig. 2a)**. This observation prompted us to investigate whether *N. benthamiana* also exhibited lethal effects on other Hemiptera insects. We conducted a similar experiment with aphids and found their mortality to be close to 100% on *N. benthamiana*, but mostly below 25% on other plants **(Fig. 2b)**. Additionally, both western flower thrips and flower thrips (Thysanoptera) experienced nearly 100% mortality on *N. benthamiana*, but only low levels of mortality (mostly below 10–15%) on other plants **(Fig. 2c, d)**. These results indicated that *N. benthamiana* possesses lethal effects on several Hemiptera and Thysanoptera pests commonly found in Solanaceae and Cucurbitaceae crops.

**Fig. 1.**
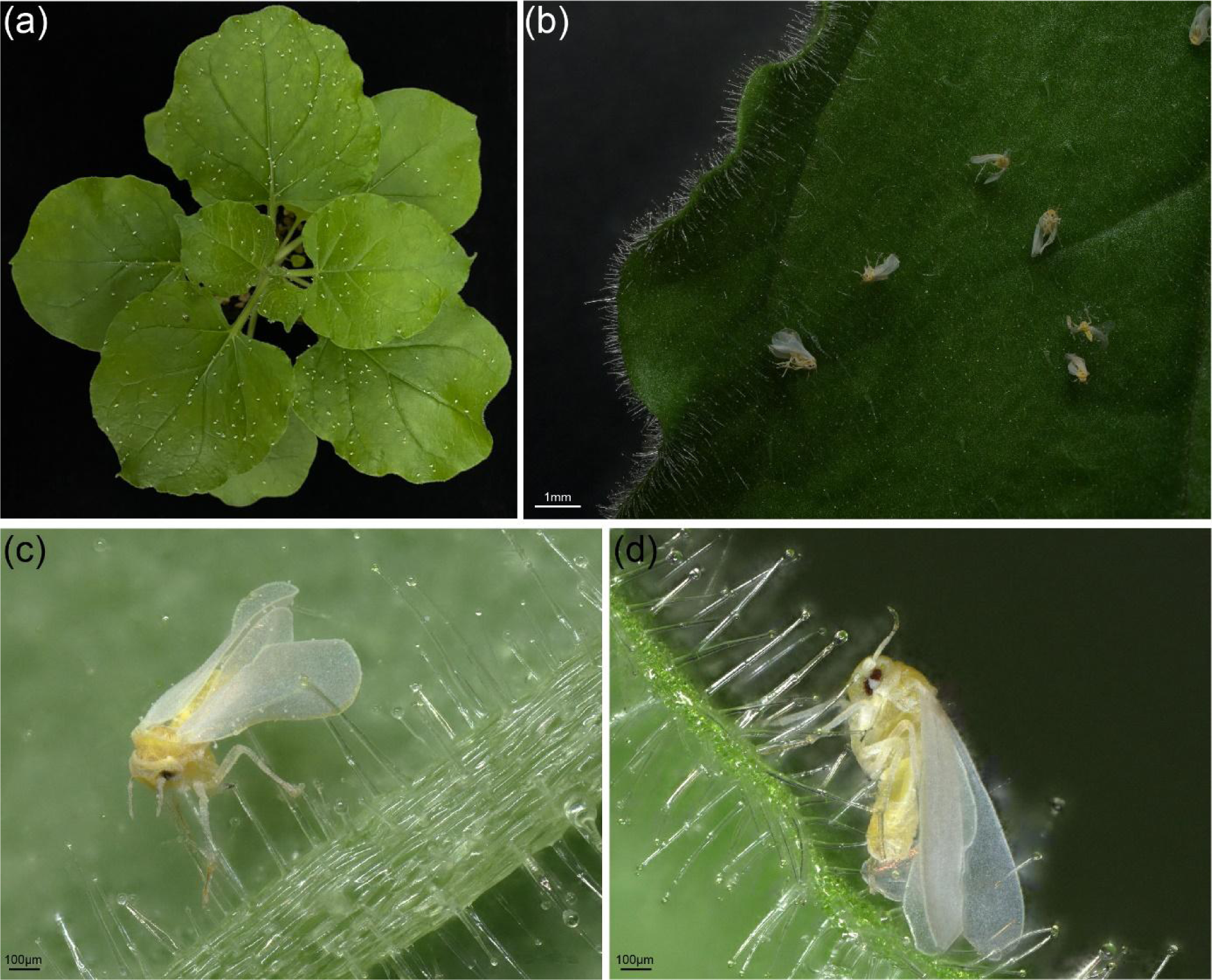
*N. benthamiana* is lethal to whiteflies. *N. benthamiana* plants display lethal effects on whiteflies, as evidenced by their dense coverage of dead whiteflies. The white dots on leaves of *N. benthamiana* are deceased whiteflies (**a**) The dead whiteflies on the *N. benthamiana* leaves show distortion and dehydration. (**b**) Dead whiteflies are notably present on the veins (**c**) and trichomes (**d**) of *N. benthamiana* plants.

**Fig. 2.**
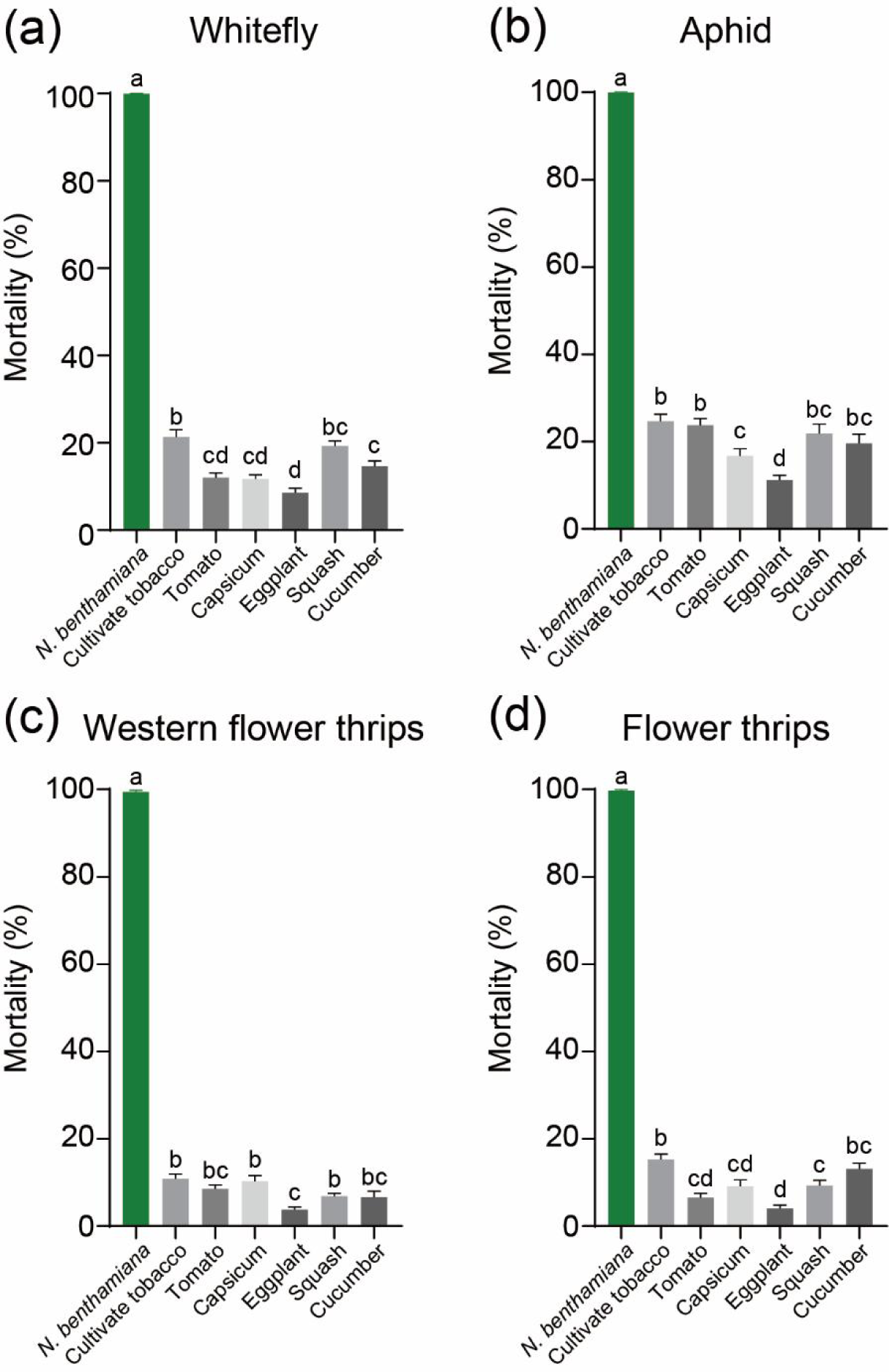
Lethal effect of *N. benthamiana* on whiteflies, aphids, and thrips in comparison to host crops. The mortality rates of whiteflies (**a**), aphids (**b**), western flower thrips (**c**), and flower thrips (**d**) were examined on both *N. benthamiana* and six crop plants (cultivated tobacco, tomato, capsicum, eggplant, squash, and cucumber). Values are mean ± SEM; *n* = 40 for **a**; *n* = 20 for **b**, **c**, and **d**. One-way ANOVA followed by Fisher’s least significant difference (LSD) test was used for significant difference analysis. Bars with different lowercase letters indicate significant differences between treatments at *p* < 0.05.

### *N. benthamiana* exhibits greater attractiveness to certain Hemiptera and Thysanoptera pests than several crops

In the greenhouse, we also observed that crop plants situated near *N. benthamiana* had fewer whiteflies and aphids compared to plants farther away. This leads us to speculate that *N. benthamiana* might be more attractive to whiteflies and aphids than other plant species. To test this hypothesis, we conducted preference tests involving the above six crop species. When given a choice between the *N. benthamiana* plant and each of the six crops, whiteflies, aphids, western flower thrips, and flower thrips consistently preferred *N. benthamiana* **(Fig. 3)**.

**Fig. 3.**
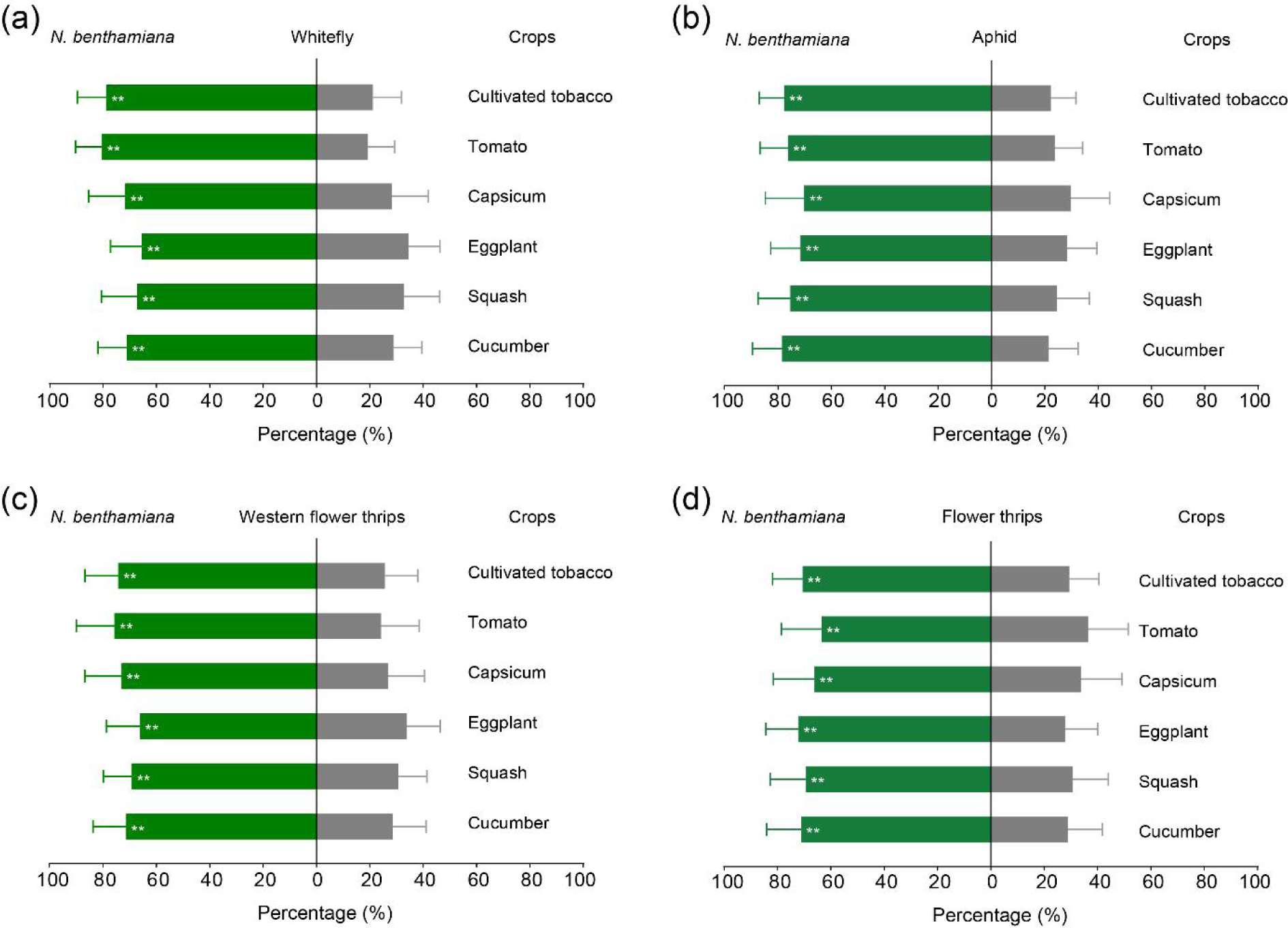
*N. benthamiana* attracts whiteflies, aphids, and thrips compared to host crops. The preference for settling on *N. benthamiana* and six crops (cultivated tobacco, tomato, capsicum, eggplant, squash, and cucumber) was assessed for whitefly (**a**), aphid (**b**), western flower thrips (**c**), and flower thrips (**d**). Values are mean ± SD, *n* = 40 for **a**; *n* = 20 for **b**, **c**, and **d**. The Nonparametric Wilcoxon matched pairs test (with two dependent samples) was used for analyzing significant differences. **, *p* < 0.01.

Plants can attract or repel insects through their shape, color, and volatile compounds. To determine if *N. benthamiana* volatiles played a role in its attractiveness to the tested Hemiptera and Thysanoptera insects, we conducted olfactometer tests. The results demonstrated that all four insects exhibited a preference for *N. benthamiana* volatiles over those emitted by the six crop species **(Fig. S5)**.

### Net cover trials using *N. benthamiana* barrier and trap whiteflies from crop tomatoes

The high attractiveness and lethal effects of *N. benthamiana* on the four Hemiptera and Thysanoptera insect pests inspired us whether it can be used as a barrier plant to trap whiteflies from nearby crops. To verify our hypothesis, two net cover trails were designed and conducted. Given that vegetable crops, such as tomatoes, are generally transplanted after seedling growth, the potential application of *N. benthamiana* during the early seedling stage was investigated.

In trial 1, we positioned potted *N. benthamiana* plants in the migration path of whiteflies to tomato plants **(Fig. S1a, b)**. Therefore, the released whiteflies would encounter the *N. benthamiana* plants first. After 24 hours, we assessed the total number of whiteflies on the tomato plants. The number of whiteflies was significantly reduced on the tomato plants located within the cover containing *N. benthamiana* along the migration route, as compared to the cover without *N. benthamiana* **(Fig. 4a)**.

**Fig. 4.**
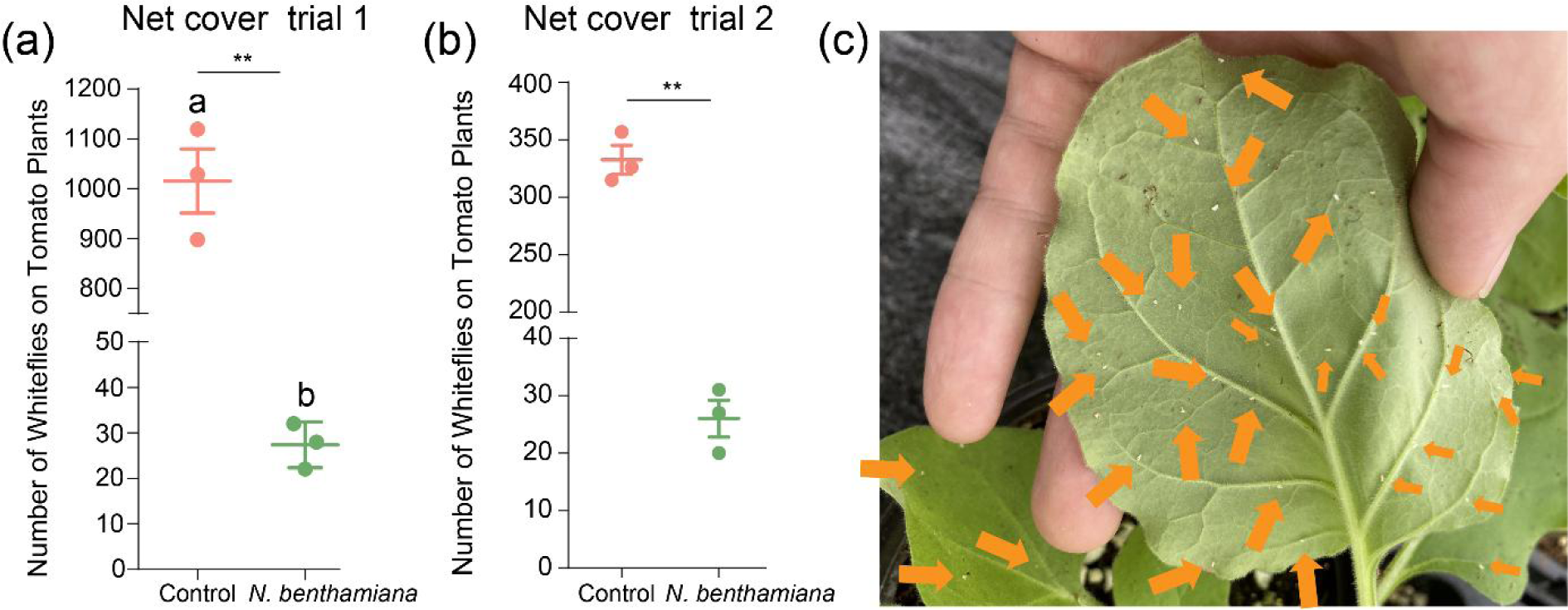
*N. benthamiana* plants barrier and trap whiteflies. Evaluation of barrier and trapping efficacy of potted *N. benthamiana* within net covers. (**a**) Result of net cover trial 1. The number of whiteflies on tomato plants: absence of potted *N. benthamiana* (Control), potted *N. benthamiana* positioned between the whitefly release point and tomato plants (Treatment: *N. benthamiana*) (italicFig. S1a, b). (**b**) Result of net cover trial 2. The number of whiteflies on tomato plants: absence of potted *N. benthamiana* (Control) or 4 potted *N. benthamiana* in the center (Treatment: *N. benthamiana*) (Fig. S1c, d). (**c**) Deceased whiteflies on potted *N. benthamiana*. Values are mean ± SEM, *n* = 3. Student’s *t*-test (two-tailed) was used for significant difference analysis. **, *p* < 0.01.

For trial 2, we assessed the attraction and lethal impact of *N. benthamiana* on whiteflies. In the net cover, *N. benthamiana* plants were circled by tomato plants and the released whiteflies would encounter tomato plants first **(Fig. S1c, d)**. Similarly, a considerable portion of the whiteflies died on the *N. benthamiana* plants situated at the center of the net cover, and only a small number of whiteflies landed on tomato plants **(Fig. 4b)**. It is noteworthy that, in both trials, the majority of the whiteflies died on the *N. benthamiana* plants **(Fig. 4c)**. These trials indicated that the introduction of *N. benthamiana* plants during the initial whitefly infestation effectively reduces their population on crops, accompanied by a pronounced lethal effect.

### Greenhouse trial 1 using *N. benthamiana* as a dead-end trap for pest control

The barrier and trap effects exhibited by *N. benthamiana* against whiteflies promote us to ascertain its potential utility as a dead-end trap plant. First, we assessed the efficacy of the field-cultivated *N. benthamiana* in controlling whiteflies. We transplanted *N. benthamiana* and tomato seedlings into an experimental field plot **(Fig. S2a-c).** In the field, the naturally occurring whiteflies in the environment will transfer to the tomato plants and the number of whiteflies increased with the growth of tomatoes **(Fig. 5a, grey dots)**. Comparatively, the number of whiteflies on the tomato crops in the plot with *N. benthamiana* plants was significantly reduced compared to the plot without them **(Fig. 5a, green dots)**. These results demonstrated that *N. benthamiana* can be employed as a dead-end trap to protect tomato crops when cultivated in the field.

**Fig. 5.**
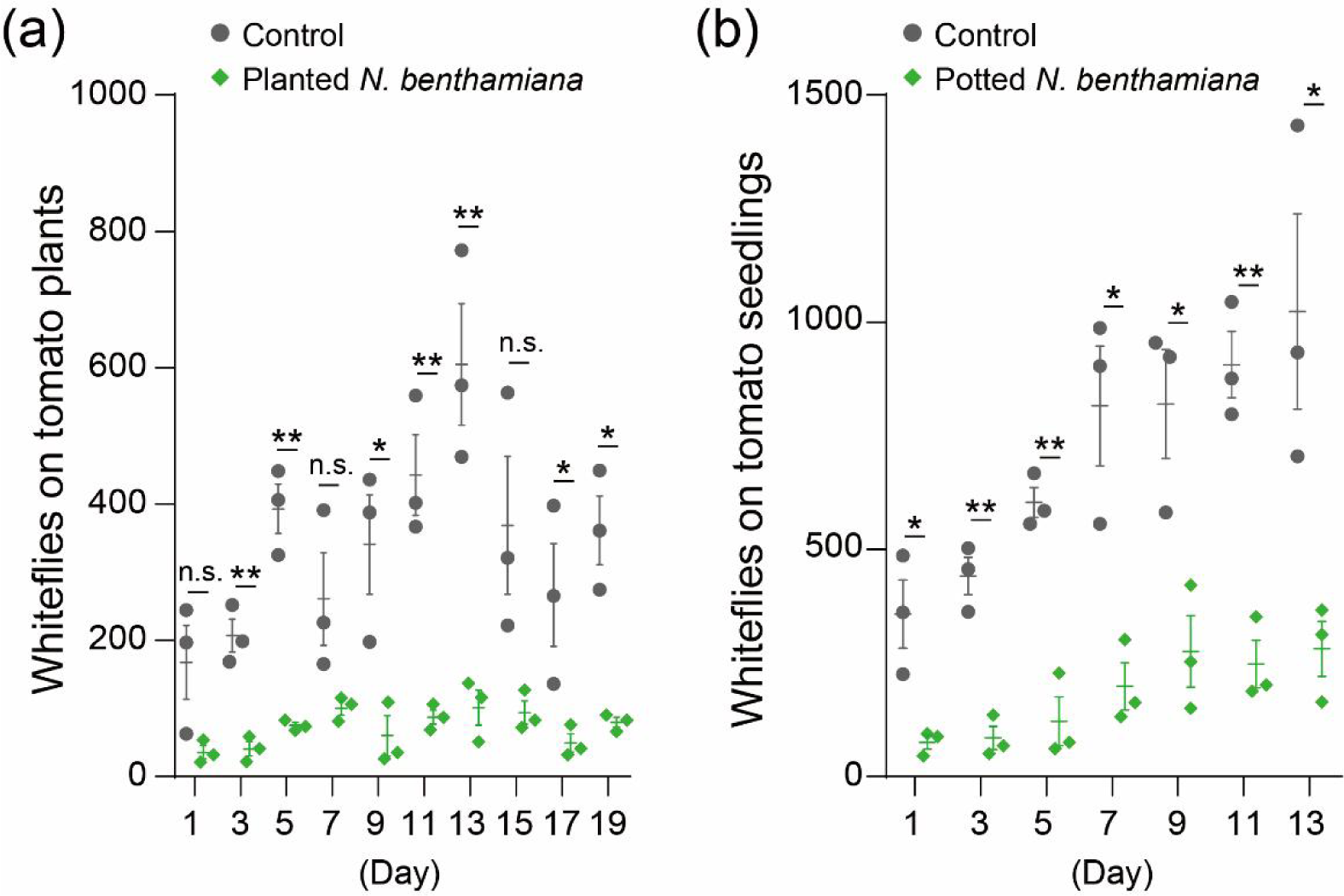
*N. benthamiana* plants as an effective dead-end trap in the greenhouse. (**a**) The transplanted *N. benthamiana* plants were utilized as dead-end traps in a greenhouse. Compared to the control plot without planted *N. benthamiana*, the plot with *N. benthamiana* transplants demonstrated a significant reduction in the number of whiteflies on the tomato. (**b**) The plot with potted *N. benthamiana* plants showed a considerable decrease in the number of whiteflies on the tomato when compared to the control plot without potted *N. benthamiana* plants. Values are mean ± SEM, *n* = 3. Student’s *t*-test (two-tailed) was used for significant difference analysis. n. s., not significant; *, *p* < 0.05; **, *p* < 0.01.

Additionally, we investigated the efficacy of potted *N. benthamiana* plants in controlling whiteflies and protecting tomato seedlings. The potted *N. benthamiana* plants were placed around the tomato seedlings **(Fig. S2d-f)**. In comparison to the plot without *N. benthamiana* pots, the number of whiteflies on the tomato seedlings in the plot with *N. benthamiana* pots was significantly reduced **(Fig. 5b)**. Notably, the *N. benthamiana* plants grew well during the experimental period in the field **(Fig. S6a)** and experimentalffectively trapped whiteflies **(Fig. S6b, comparison)**. These findings confirmed that both field-cultivated and potted *N. benthamiana* can serve as effective dead-end traps for protecting tomato crops against whiteflies in the field.

### Greenhouse trial 2 using *N. benthamiana* as a substitute for commercial sticky traps

The current method of trapping Hemiptera and Thysanoptera pests involves the use of artificial plastic sticky traps (Steiner *et al*., 1999). However, the viscosity of the commercial traps decreases in the field, and their usage contributes to plastic pollution in the environment. In light of this, we evaluated whether the *N. benthamiana* plants could serve as a viable substitute for commercial sticky traps in attracting and killing pests naturally occurring in the experimental greenhouse. The experimental tomato greenhouse was divided into four plots, where potted *N. benthamiana* plants and yellow or blue sticky traps were placed **(Fig. S3)**.

Subsequently, we counted the number of naturally occurring whiteflies and thrips on the *N. benthamiana* plants, sticky traps, and main crops. Our observations revealed that both the *N. benthamiana* plants and the yellow sticky traps attracted a significant number of whiteflies and thrips, while fewer were attracted to the blue sticky traps **(Fig. 6a, b)**. In relation to this, it was noted that the population of whiteflies, but not thrips, was significantly reduced on tomatoes sheltered by *N. benthamiana* plants when compared to tomatoes within the control plots **(Fig. 6c, d)**. These results indicated that the trapping efficacy of the *N. benthamiana* plants for naturally occurring whiteflies and thrips was comparable to that of yellow sticky traps and superior to that of blue traps.

**Fig. 6.**
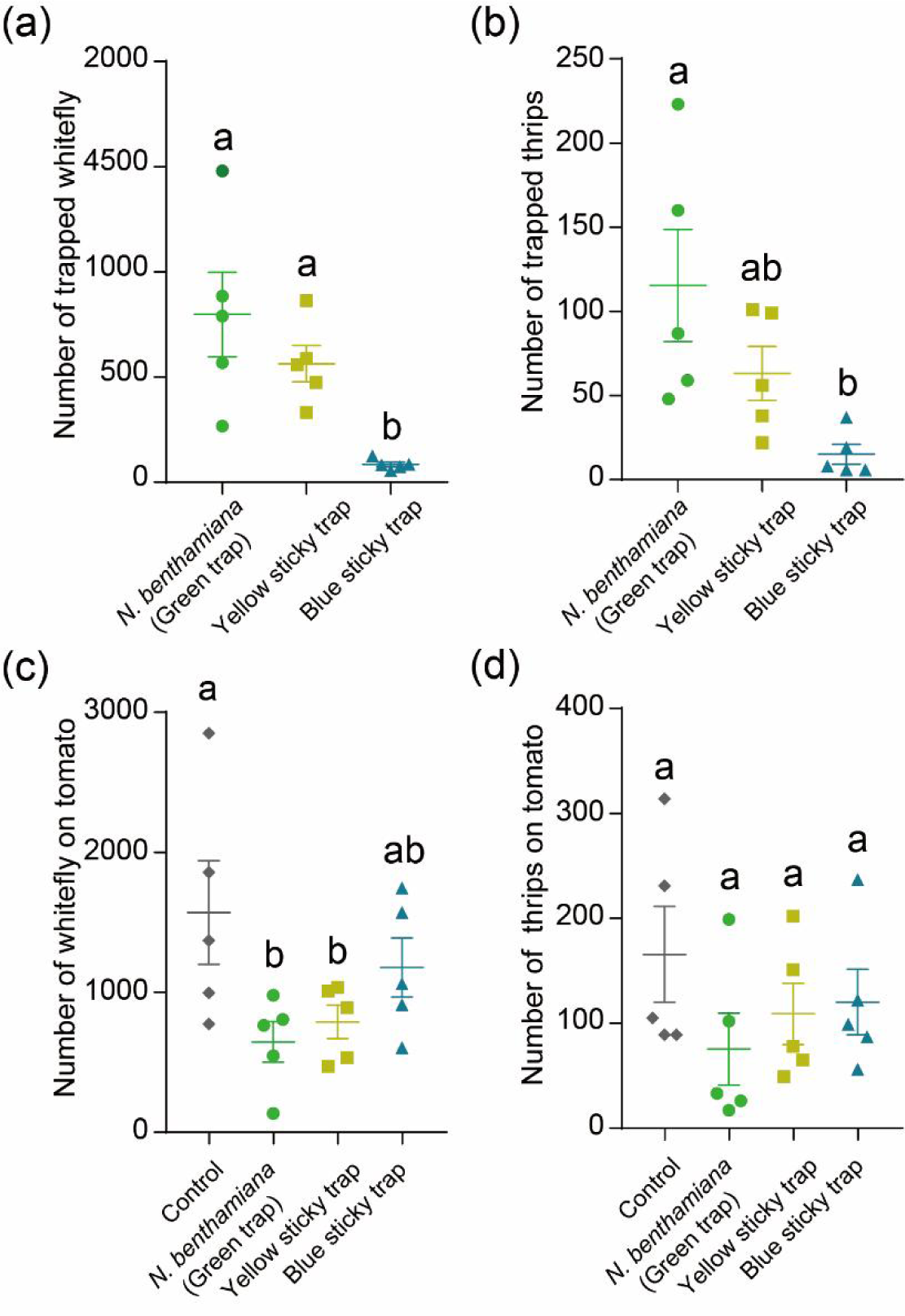
*N. benthamiana* plants as a substitute for commercial sticky traps. The number of trapped whiteflies (**a**) and thrips (**b**) on *N. benthamiana*, yellow sticky traps, and blue sticky traps. The numbers of whiteflies (**c**) and thrips (**d**) on tomato plants were assessed in control and three treatment conditions: the *N. benthamiana* plant, yellow sticky trap, and blue sticky trap. Values are mean ± SEM, *n* = 5. One-way ANOVA followed by Fisher’s least significant difference (LSD) test was used for significant difference analysis. Bars with different lowercase letters indicate significant differences between treatments at *p* < 0.05.

To further assess the pest control effect, we evaluated the efficacy of the *N. benthamiana* plant and commercial sticky traps in controlling pests on pepper and cucumber plants in the greenhouse. Similarly, we found that the *N. benthamiana* plant trapped more whiteflies and thrips compared to the commercial blue sticky traps, leading to a reduction in pest numbers on pepper **(Figs. S7)** and cucumber plants **(Figs. S8)**. These findings demonstrated that *N. benthamiana* plants can serve as effective dead-end traps in the greenhouse for controlling whitefly and thrip pests on both Solanaceae and Cucurbitaceae crops. Moreover, they have the potential to serve as an environmentally friendly alternative to commercial plastic sticky traps.

### Commercial greenhouse trials of *N. benthamiana* as a dead-end trap for pest control

To further evaluate the effectiveness of the *N. benthamiana* plant as a dead-end trap for field pest management, we conducted trials in a commercial tomato greenhouse **(Fig. S4a, b).** The greenhouse was divided into five experimental plots, where we evaluated and compared the use of potted and field-transplanted *N. benthamiana* plants with commercial sticky blue and yellow traps, as well as a control group **(Fig. S4c)**. Main crop tomato plants were raised in seedling trays and then transplanted into the commercial greenhouse. Simultaneously, *N. benthamiana* plants were transplanted **(fig. S4d)**, and potted plants **(Fig. S4e),** and commercial sticky traps were placed in different plots.

The number of whiteflies on tomato crops in the five experimental plots was continuously monitored every seven days from September 3^rd^ to October 29^th^, 2022. In the experimental area, only whitefly and a small number of larvae of *Spodoptera litura* occurred naturally druing our trial. Similar to the greenhouse trials mentioned above, both transplanted and potted *N. benthamiana* plants grew well in the commercial greenhouse during the entire experimental period **(Fig. S9)**. We found that the number of whiteflies on tomatoes in the control and experimental plot with the blue sticky trap was significantly higher than in the plot with the yellow sticky trap, potted *N. benthamiana* plants, and field-transplanted *N. benthamiana* plants **(Fig. 7)**. In the early stages of the trial, the yellow sticky trap demonstrated slightly better control effect than the *N. benthamiana* plants (observed on September 3^rd^ and 10^th^ in Fig. 7). However, in the middle and late stages, the *N. benthamiana* plants exhibited better control, likely due to their continuous growth (observed on October 1^st a^nd 22^nd i^n Fig. 7). These results demonstrated that both potting and field planting of *N. benthamiana* plants effectively controlled whiteflies, and showed comparable or superior control compared to commercial sticky traps.

**Fig. 7.**
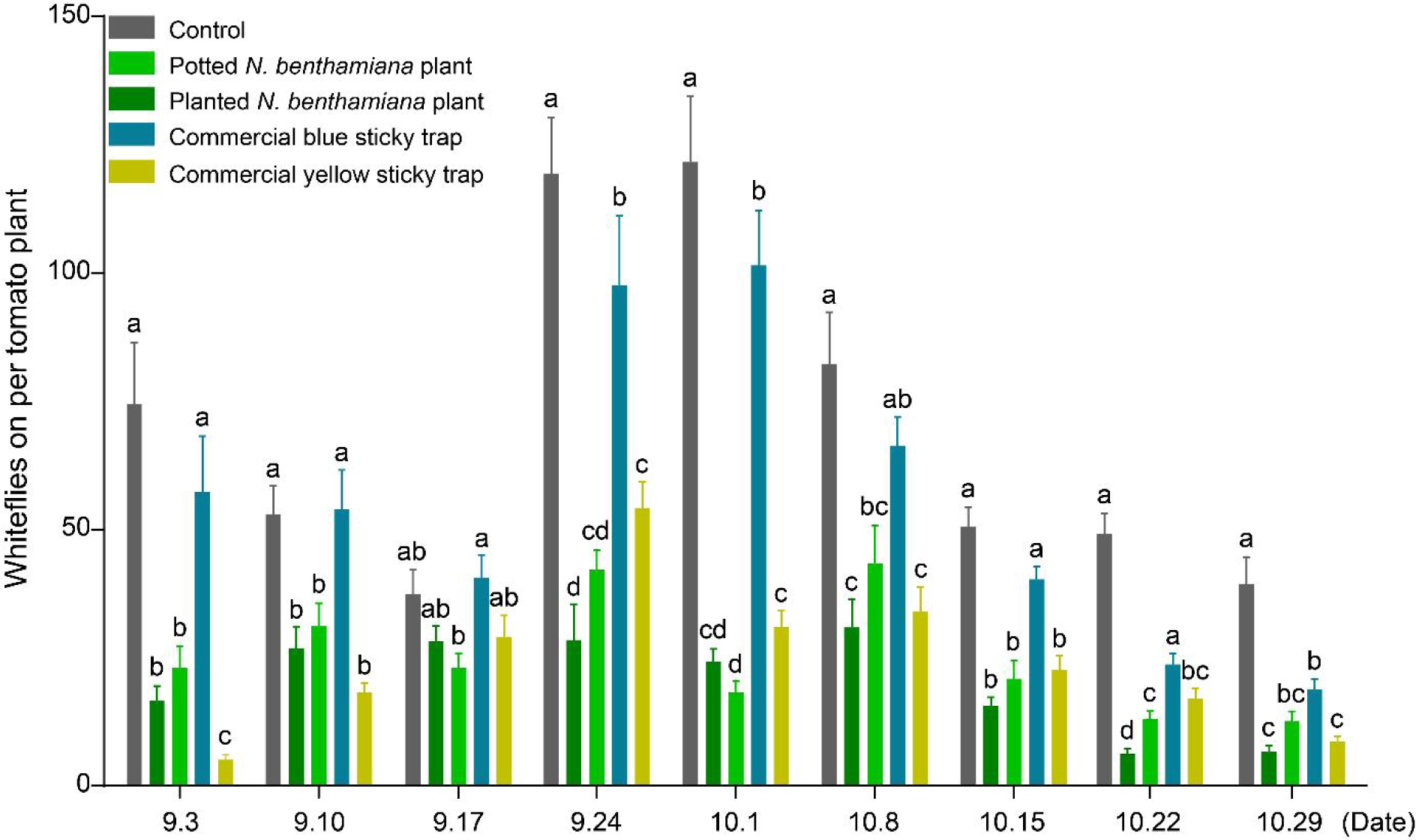
*N. benthamiana* plants for pest control in a commercial greenhouse. In a commercial greenhouse trial, the number of whiteflies on the main crop tomatoes was compared across experimental plots with potted or planted *N. benthamiana* plants, commercial yellow or blue sticky traps, and control plots. Values are mean ± SEM, *n* = 24. One-way ANOVA followed by Fisher’s least significant difference (LSD) test was used for significant difference analysis. Bars with different lowercase letters indicate significant differences between treatments at *p* < 0.05.

### *N. benthamiana* is a crop plant and natural enemy-friendly trap plant

The use of trap crops in the field can potentially interfere with crop growth through nutrient competition or light obstruction. To assess whether the introduction of *N. benthamiana* plants in the field had any negative effects on the main crops, we also evaluated the growth of tomatoes during the trials. The results indicated that, for tomatoes, the early branching number, late flowering number, and individual height of tomato plants were not significantly impacted by the presence of *N. benthamiana* **(Fig. S10)**.

Commercial sticky boards used for trapping pests also capture and kill beneficial insects and natural enemies, rendering them incompatible with measures involving the application of natural enemies. In our trials, we observed that yellow sticky traps cause significant mortality to Hymenoptera insects and natural enemies of whiteflies including the predatory natural enemy *Nesidiocoris tenuis* and the parasitism natural enemy wasp *Eretmocerus hayati*. However, no such natural enemies were observed on the *N. benthamiana* plants. Further laboratory experiments showed that the *N. benthamiana* plant had no lethal effect on *Nesidiocoris tenuis* and honeybees (*Apis mellifera*) **(Fig. S11)**. These findings indicated that the use of the *N. benthamiana* plant as a dead-end trap does not harm these natural enemies and pollinators, providing an advantage over the application of conventional sticky traps.

## Discussion

The use of chemical pesticides remains a major concern in agriculture, ecology, and society. Therefore, the search for safe and environmentally benign alternatives is of utmost importance. Hemiptera and Thysanoptera insect pests pose significant threats to various important crops, and are notorious for their rapid development of pesticide resistance, making them difficult to control. In this study, we conducted net cover trials and three field assays over two years to demonstrate the effectiveness of the laboratory model plant *N. benthamiana* as a dead-end trap for Hemiptera and Thysanoptera pests **(Figs. 3–7)**. Our findings support the use of *N. benthamiana* as a substitute for commercial sticky traps, controlling pests while reducing the environmental impact associated with plastic pollutants. Furthermore, the implementation of *N. benthamiana* traps does not affect the natural enemies of these pests **(Fig. S11)**, making italict conducive to integrated IPM strategies. Our work introduces a novel application for the well-known *N. benthamiana* plant in the greenhouse of pest control, providing an easily applicable and sustainable measure for managing Hemiptera and Thysanoptera pests **(Fig. 8)**.

**Fig. 8.**
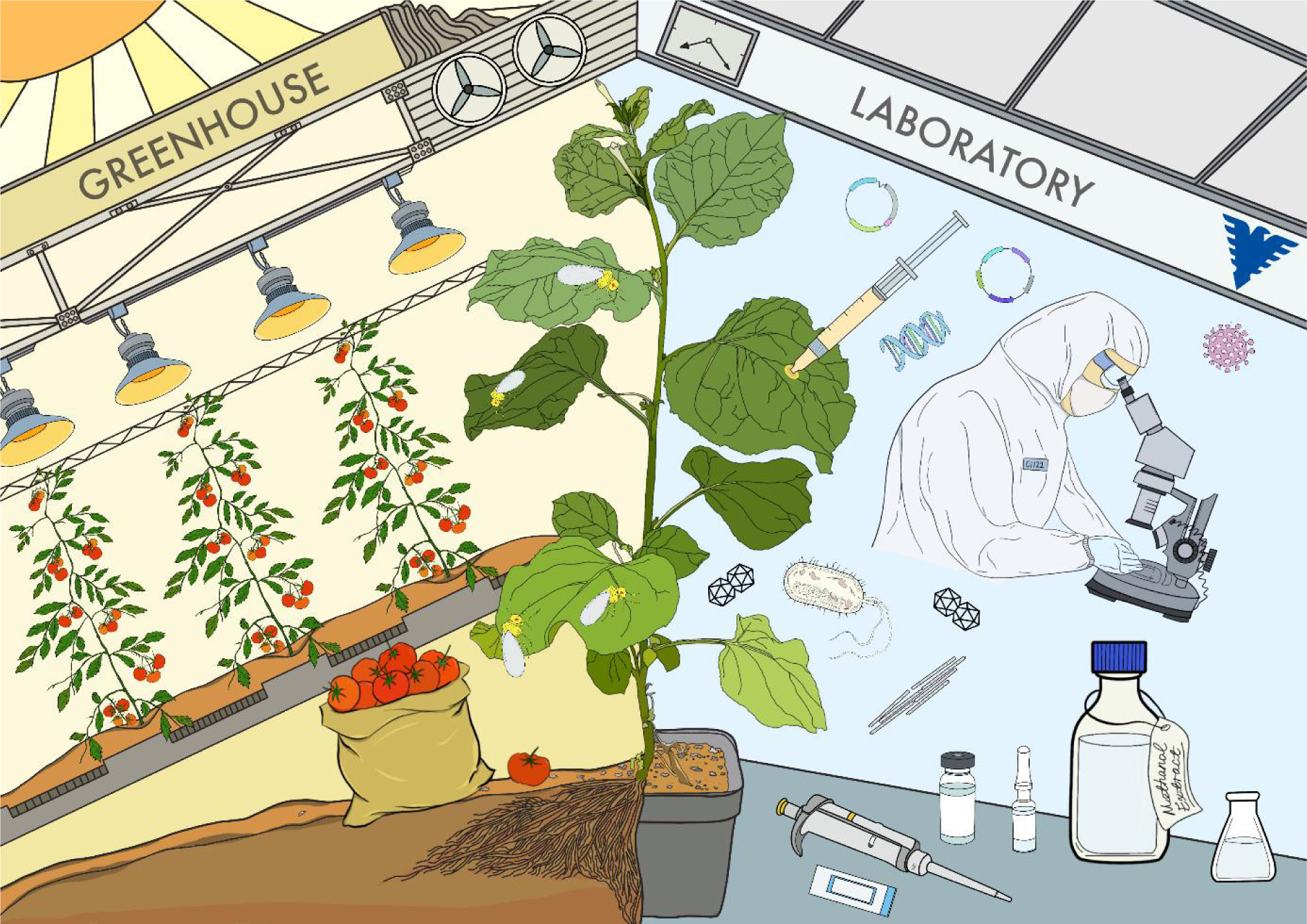
New value of model plant *N. benthamiana*: From laboratory to field, from model plant to dead-end trap. Right, *N. benthamiana*, a classical lab model plant, is extensively employed in scientific research, spanning biopharmaceutical applications (for example, vaccine production) and playing a pivotal role in uncovering RNA interference (RNAi), plant-pathogen interactions (plant fungi, bacteria, and viruses), metabolic pathway engineering, synthetic biology, and gene editing. Left, the less common ecological interactions between *N. benthamiana* and insects. *N. benthamiana* has potent allure and lethal impact on several Hemiptera and Thysanoptera species. As a model plant, *N. benthamiana* has the potential to serve as a dead-end trap in crop and vegetable cultivation.

To the best of our knowledge, few dead-end trap plants specifically targeting Hemiptera and Thysanoptera pests has been reported previously. Previous studies have shown that certain trap plants, such as vetiver grass *Vetiveria zizanioides* for controlling rice striped stem borer *Chilo suppressalis* (Lu *et al*., 2019) and *Barbarea vulgaris* for controlling *Plutella xylostella* (Shelton & Nault, 2004; Badenes-Perez *et al*., 2005a), can effectively attract adult females to lay eggs but prevent larvae from completing their life cycle. In contrast, our study reveals that *N. benthamiana* plants not only attract whiteflies and thrips but also kill these pests at the adult stage **(Figs. 1–3)**. *N. benthamiana* plants can serve as dead-end traps when cultivated in the field or used as potted plants, both methods showing promising pest control effects **(Figs. 4–7)**. Field-cultivated *N. benthamiana* plants can be planted alongside crops, while potted *N. benthamiana* plants offer flexibility and can serve as a substitute for commercial sticky traps. By adapting the placement of these plants according to pest occurrence in greenhouses or other facilities, more effective pest control can be achieved.

The selection of main crops is a crucial factor to consider when employing dead-end traps in the field. Main crops should be chosen based on their lower attractiveness to pests compare to trap plants, taking into account the specific target pests. Previous reports on dead-end trap plants have focused on attracting a single insect species and have been limited to specific main crop species (Thompson, 1988; Idris & Grafius, 1996; Lu *et al*., 2004; Shelton & Badenes-Perez, 2006). In our study, we found that all tested Solanaceae and Cucurbitaceae crops were less attractive to Hemiptera and Thysanoptera pests compared to *N. benthamiana* plants **(Figs. 3, S5)**. Greenhouse trials demonstrated that *N. benthamiana* plants effectively trapped pests on tomatoes, peppers, and cucumbers **(Figs. 4–7, S7, S8)**. Our results suggested that *N. benthamiana* plants may serve as dead-end trap plants for a wide range of crop species, enabling simultaneous control of multiple pests.

The adaptability of dead-end trap plants in crop fields significantly affects their practical application. *N. benthamiana* plants are originally from the desert in Central Australia, characterized by harsh climatic conditions (Bally *et al*., 2018). Despite *N. benthamiana*’s exclusive Australian natural distribution, our field trials demonstrated its adaptability to the monsoon climatic region of southern China. The ability of *N. benthamiana* to thrive in extreme and variable environments further underscores its potential as a dead-end trap for plant pest control. While *N. benthamiana* is considered pathogen-sensitive, both open-field and commercial greenhouse trials revealed normal plant growth, with *N. benthamiana* plant being less severely affected or killed by various pathogens than anticipated **(Figs. S6a, S9)**. This further validates the viability of *N. benthamiana* as a dead-end trap for field pest control.

The use of sticky traps increases the cost of pest management and contributes to plastic pollution in the environment. Additionally, the application of sticky traps in the field can harm beneficial insects, including pollinators and natural enemies, hindering their joint use in pest control. Our greenhouse trials demonstrated that *N. benthamiana* plants trap an equivalent number of whiteflies or thrips compared to yellow sticky boards, effectively reducing pests on main crops **(Figs. 6, S7, S8)**. Furthermore, no Hymenoptera insects or natural enemy *N. tenuis* were found on *N. benthamiana* plants, and laboratory tests confirmed that *N. benthamiana* plants had no lethal effect on honeybees and *N. tenuis* **(Fig. S11)**. Moreover, while sticky traps require regular replacement to maintain effectiveness, *N. benthamiana* plants can continue to grow for 5–6 months, effectively trapping pests throughout this period. Certain trap plants may have adverse effects on the main crops, such as depleting soil nutrients originally intended for crops, blocking light, or even secreting allelochemicals that affect crop growth. In our commercial greenhouse trials, field-grown *N. benthamiana* plants showed no significant impact on the early growth, branching, and later growth and flowering of main crop tomatoes **(Fig. S10)**. These resultsts suggested that *N. benthamiana* plants may serve as an ideal substitute for plastic sticky traps in IPM approaches.

In this study, we reported the remarkable resistance and attractiveness of the laboratory model plant *N. benthamiana* to Hemiptera and Thysanoptera insects. Our observations indicate that trichomes on *N. benthamiana* probabaly trap and kill whiteflies **(Fig. 1b–d).** We speculate that these trichomes may secrete whitefly-toxic chemicals. Further exploration into the underlying mechanisms of *N. benthamiana*’s resistance against Hemiptera and Thysanoptera pests is both justified and essential. In addition, several pest species showed a preference for *N. benthamiana* volatiles, and further work can identify components of these volatiles that attract pests **(Figs. 2, 3, S5)**.

Based on these characteristics of *N. benthamiana*, we proposed that it can serve as a green trap for field pest control, offering an alternative to commercial sticky traps. It effectively controls pests throughout the entire crop growth period. Meanwhile, the field *N. benthamiana* does not affect the growth of the main crop and the pest’s natural enemies, which is an ideal strategy for IPM. Overall, our work unveils a previously undiscovered attribute of the prominent laboratory model plant *N. benthamiana*, underscoring its potential to play a vital role in sustainable field pest control **(Fig. 8)**.

## Acknowledgements

Financial support for this paper was provided by the National Key Research and Development Program (2021YFC2600100), the National Natural Science Foundation of China (31925033), Tobacco Pests and Diseases Green Prevention and Control Major Special Project (110202101045, LS-05), and the earmarked fund for China Agriculture Research System (CARS-23-C05). We thank Ms. Rong Jin from Zhejiang University for greenhouse management.

## Author contributions

W-HH, J-XW, and X-WW conceived and designed the study. W-HH, J-XW, F-BZ, S-XJ, Y-WZ, and Y-QL performed the experiments. W-HH, J-XW, and F-BZ analyzed the data. W-HH, J-XW, F-BZ, and X-WW wrote and finalized the manuscript.

## Data availability

**Supplementary Information** is available for this paper. Correspondence and requests for materials should be addressed to Xiao-Wei Wang or Wen-Hao Han.

## Supporting Information

**Fig. S1.** The layout of the experimental plots of the net cover trials.

**Fig. S2.** The layout of greenhouse trial 1 for eluting *N. benthamiana* as a dead-end trap.

**Fig. S3.** The layout of pepper, cucumber, and tomato in greenhouse trial 2 to compare the effect between *N. benthamiana* and commercial sticky traps.

**Fig. S4.** The layout of the commercial greenhouse trial.

**Fig. S5.** Preference insect pests in olfactometer for *N. benthamiana* and several crops.

**Fig. S6.** Field-grown *N. benthamiana* plants attract and kill whiteflies.

**Fig. S7.** *N. benthamiana* plants as a substitute for commercial sticky traps in pepper greenhouse.

**Fig. S8.** *N. benthamiana* plants as a substitute for commercial sticky traps in cucumber greenhouse.

**Fig. S9.** The status of field-grown *N. benthamiana* plants in the commercial greenhouse.

**Fig. S10.** Traits of the main crop tomatoes in experimental plots with *N. benthamiana* plants and sticky traps.

**Fig. S11.** Effects of *N. benthamiana* plants on beneficial insects.

