## Supporting Information for "A New Feature of the Laboratory Model Plant *Nicotiana benthamiana*: Dead-End Trap for Sustainable Field Pest Control"

The following Supporting Information is available for this article:

**Fig. S1.** The layout of the experimental plots of the net cover trials.

**Fig. S2.** The layout of greenhouse trial 1 for eluting *N. benthamiana* as a dead-end trap.

**Fig. S3.** The layout of pepper, cucumber, and tomato in greenhouse trial 2 to compare the effect between *N. benthamiana* and commercial sticky traps.

**Fig. S4.** The layout of the commercial greenhouse trial**.**

**Fig. S5.** Preference of insect pests in olfactometer for *N. benthamiana* and several crops.

**Fig. S6.** Field-grown *N. benthamiana* plants attract and kill whiteflies.

**Fig. S7.** *N. benthamiana* plants as a substitute for commercial sticky traps in pepper greenhouse.

**Fig. S8.** *N. benthamiana* plants as a substitute for commercial sticky traps in cucumber greenhouse.

**Fig. S9.** The status of field-grown *N. benthamiana* plants in the commercial greenhouse.

**Fig. S10.** Traits of the main crop tomatoes in experimental plots with *N. benthamiana* plants and sticky traps.

**Fig. S11.** Effects of *N. benthamiana* plants on beneficial insects.

### Supplementary Figures and Legends

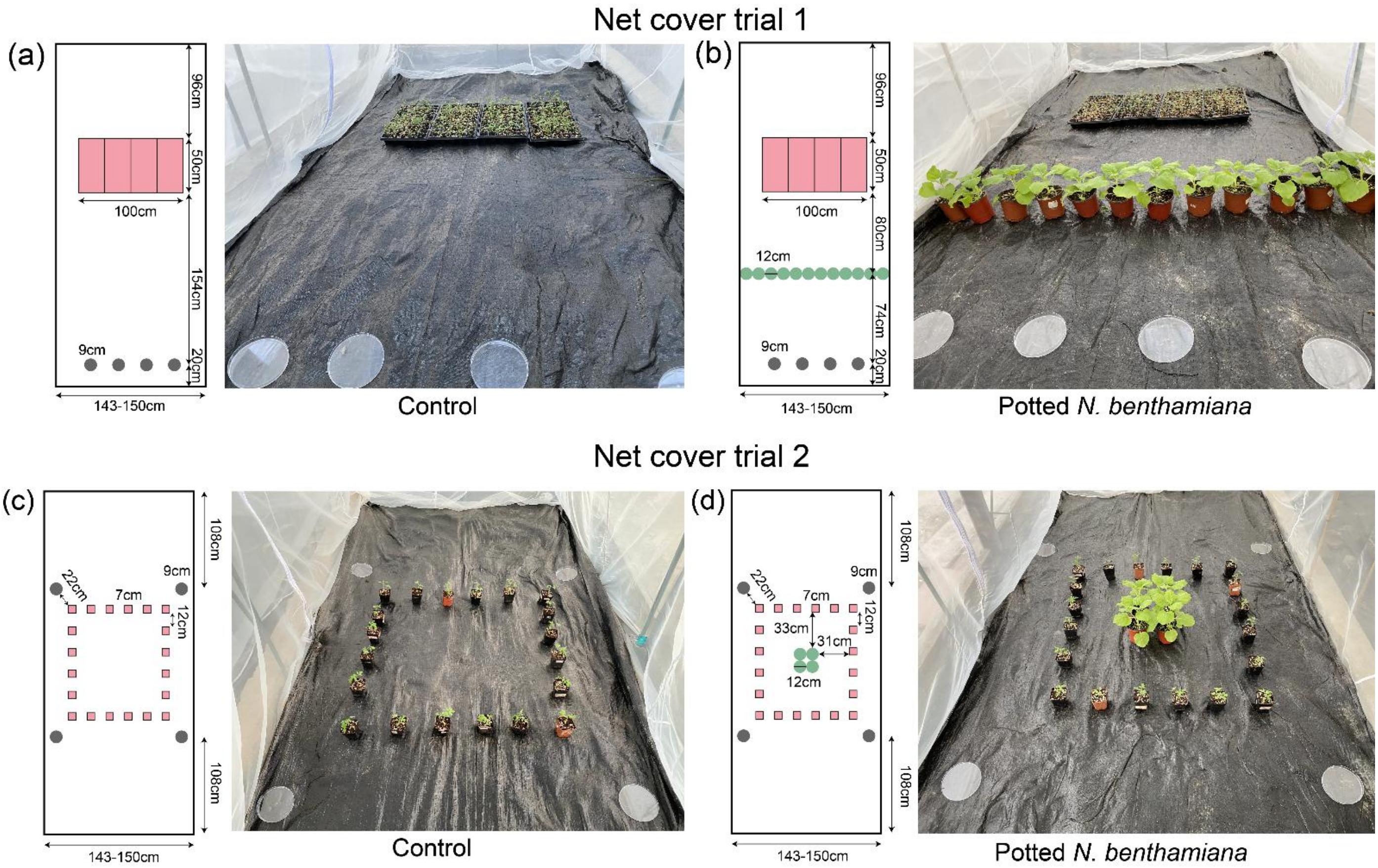


**Fig. S1. The layout of the experimental plots of the net cover trials.**

The layout of net cover trials. **a**, and **b** for net cover trial 1. Experimental plots **a** (Control), and **b** (*Potted N. benthamiana* plants positioned between whitefly release points and tomato plants) were designed to evaluate the barrier effect of potted *N. benthamiana* plants on whiteflies. **c** and **d** for net cover trial 2. Experimental plots **c** (Control) and **d** (Potted *N. benthamiana* plants encircled by tomato plants were designed to evaluate the trapping effect of potted *N. benthamiana* plants on whiteflies. Each diagram includes both the top-view parameters and corresponding actual photographs. In all diagrams, pink rectangles indicate a 50-hole seedling tray, pink squares indicate potted tomato plants, dark green circles indicated potted *N. benthamiana* plants, and grey circles indicated the whitefly release point where the culture dish was placed.

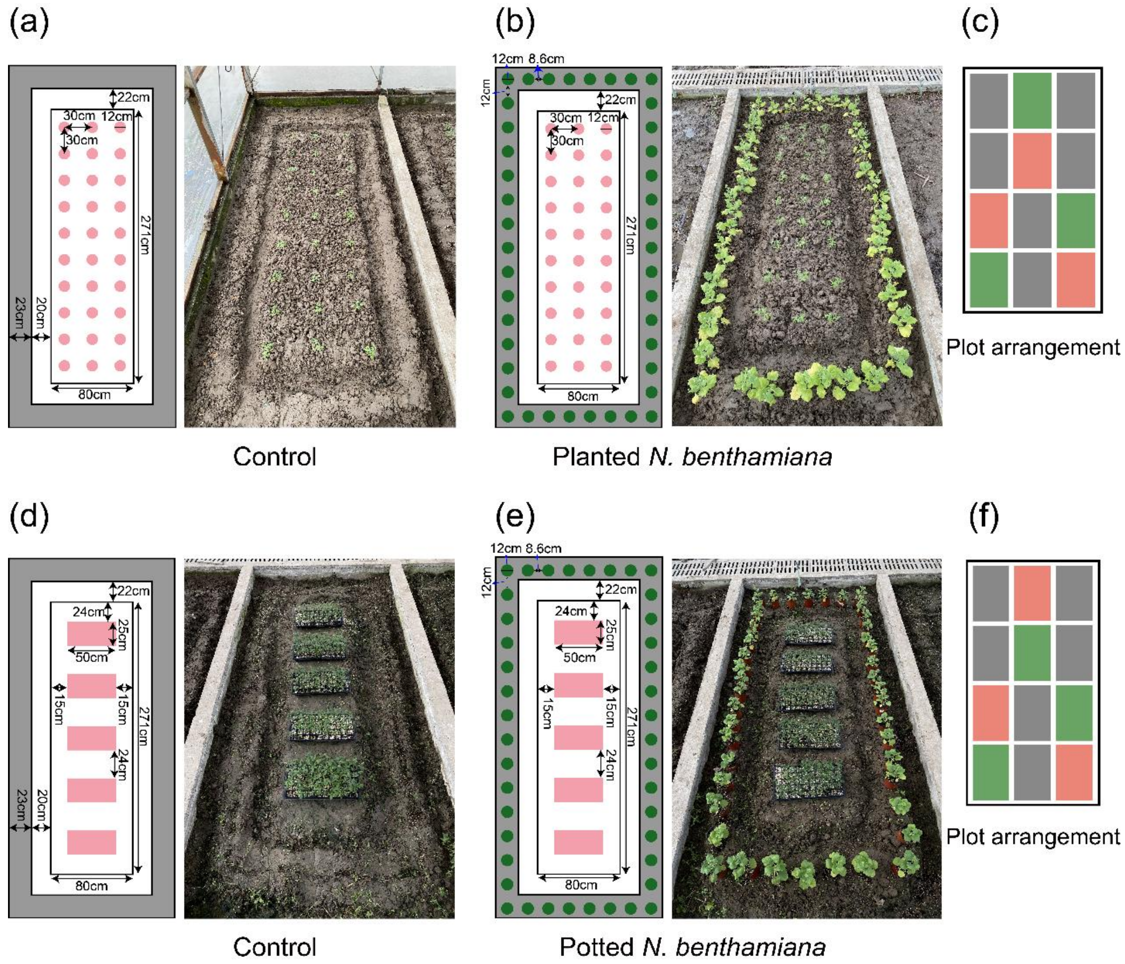


**Fig. S2. The layout of greenhouse trial 1 for eluting *N. benthamiana* as a dead-end trap.**

The design of Greenhouse trial 1 encompassed the layout of the experimental plots. Two distinct scenarios were investigated: Experimental plots **a** (Control) and **b** (Planted *N. benthamiana*). These plots were desvised to evaluate the efficacy of field-planted *N. benthamiana* as a dead-end trap plant for whitefly control. In diagrams A and B, pink circles indicate tomato seedlings cultivated within the field, while dark green circles indicate planted *N. benthamiana*. (**c**) The plot configuration encompassed 12 plots within the experimental greenhouse: three allocated to the control plot (pink), three to the treatment plot (green), and six to the non-set plot (gray). Experimental plots **d** (Control) and **e** (Potted *N. benthamiana* plants) as designed to assess the efficacy of potted *N. benthamiana* plants as a dead-end trap for whitefly control. Diagrams **d** and **e** feature pink rectangles indicating tomato seedlings within nursery trays, and light green circles indicating potted *N. benthamiana* plants. The similarly plot arrangement **f**. The experimental greenhouse is encompassed by the natural environment, facilitating the transfer of whiteflies from the surrounding environment to the plants within the experimental setup.

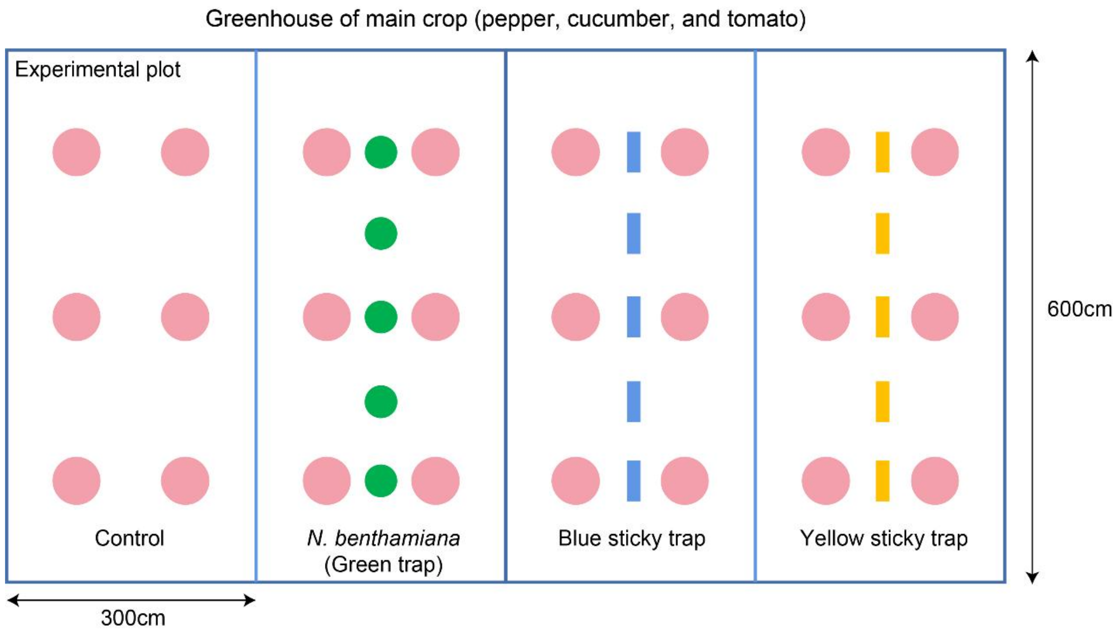


**Fig. S3. The layout of pepper, cucumber, and tomato in greenhouse trial 2 to compare the effect between *N. benthamiana* and commercial sticky traps.**

The experimental arrangement succinctly illustrates the spatial relationship among the main crops, *N. benthamiana* plants, and commercial sticky traps in greenhouse trial 2. Each greenhouse cubicle was subdivided into dual plots via a mesh grid boasting a 120-mesh count. Within each of these plots, an assemblage of six tomato, pepper, or cucumber plants was cultivated. Four partitioned plots (each comprising two separate cubicles) were established. Within these plots, potted *N. benthamiana* plants were placed, accompanied by either commercial yellow or blue sticky traps. This constituted an individual experimental trial, and this process was repeated five times, generating five independent replicates for every primary crop species. Pink circles indicate the main crops tomato, pepper, or cucumber; Green circles indicate the potted *N. benthamiana* plants; And blue or yellow rectangles indicate blue or yellow sticky traps.

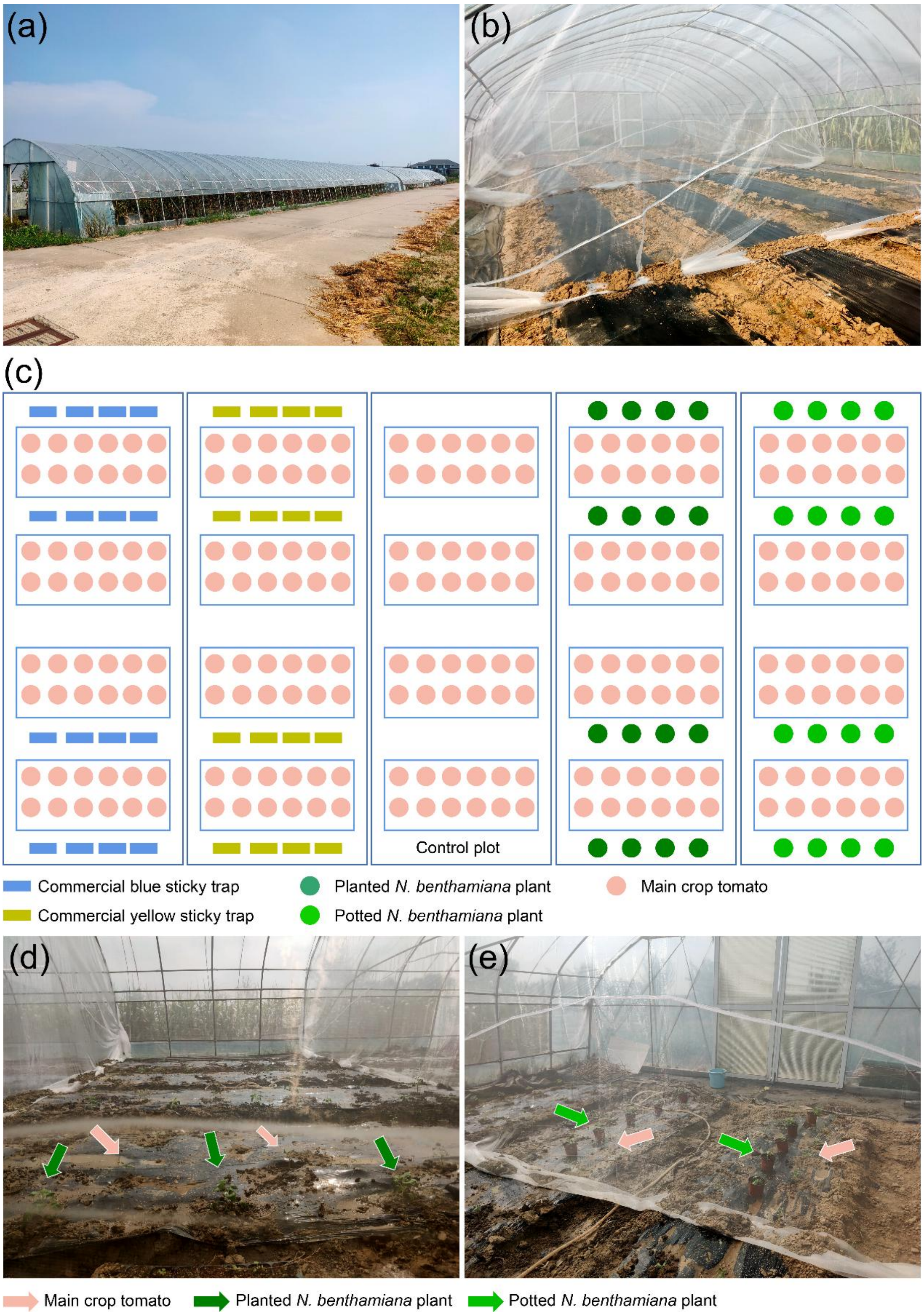


**Fig. S4. The layout of the commercial greenhouse trial.**

The commercial greenhouse for experiments was divided into five experimental plots using insect-proof nets (**a, b**). The bottom of the nets near the ground is buried in the field when the trial was conducted. The diagram of the commercial greenhouse trial (**c**), with the plots containing planted (**d**) and potted (**e**) *N. benthamiana* plants.

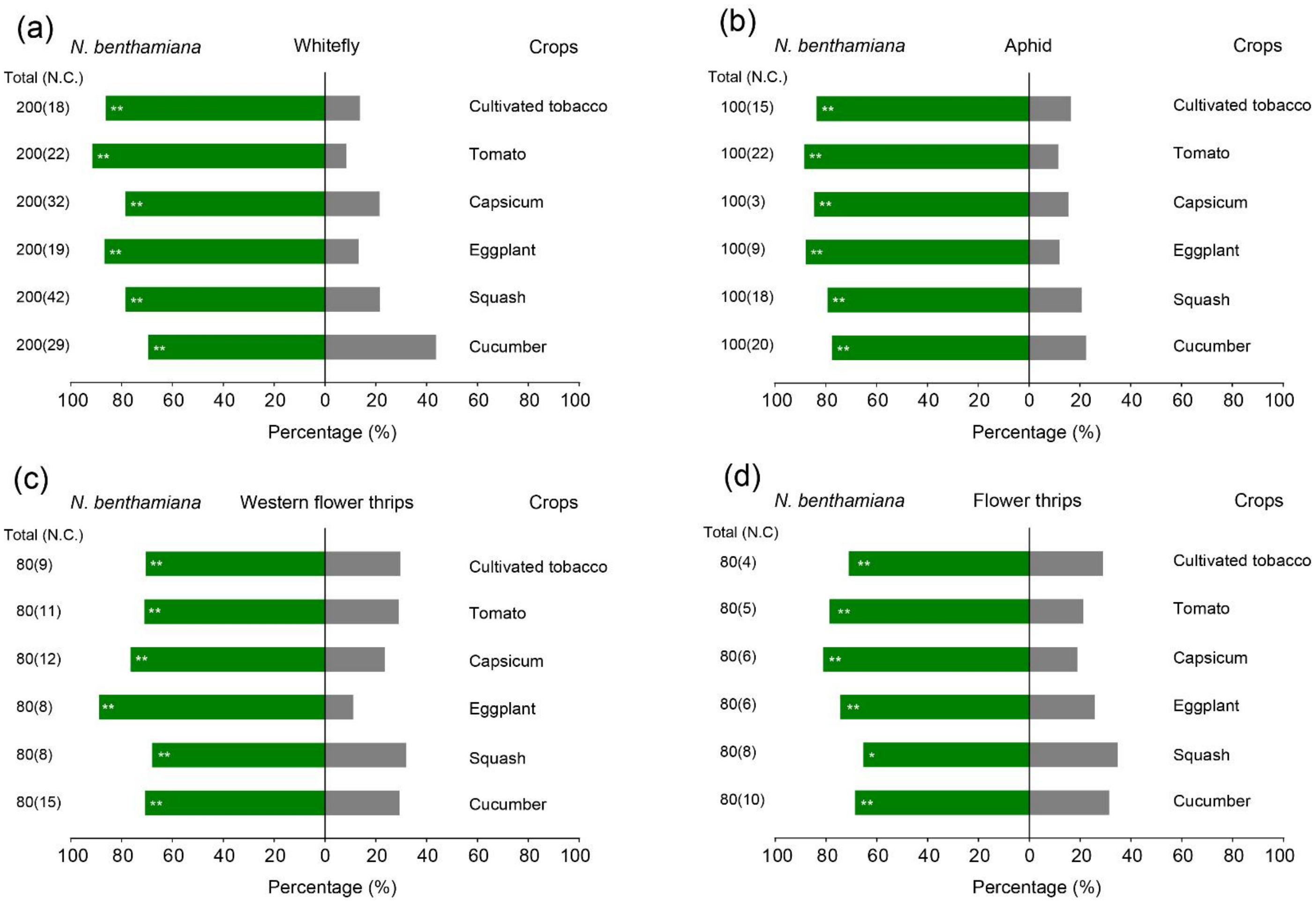


**Fig. S5. Preference of insect pests in olfactometer for *N. benthamiana* and several crops.**

The olfactometer preference of whiteflies (**a**), aphids (**b**), western flower thrips (**c**), and flower thrips (**d**) for *N. benthamiana* and six crops (cultivated tobacco, tomato, capsicum, eggplant, squash, and cucumber). "Total (N.C.)" indicates the total number of tested insects (number of insects showing no choice). The binomial distribution test was utilized for analyzing significant difference analysis. *, $p < 0.05$. **, $p < 0.01$.

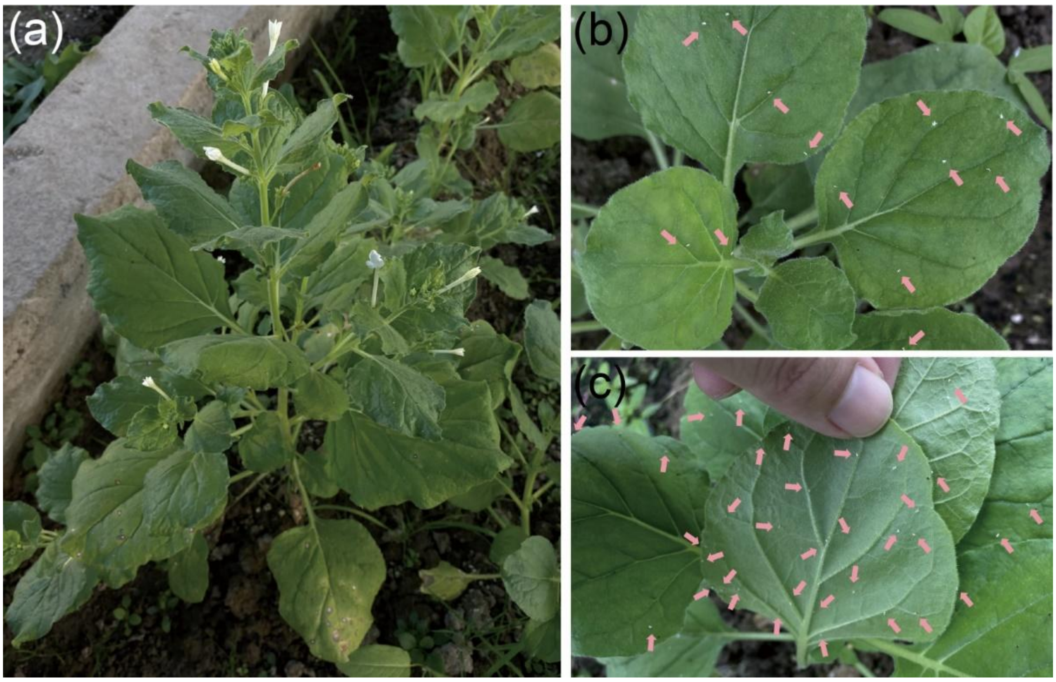


**Fig. S6. Field-grown *N. benthamiana* plants attract and kill whiteflies.**

(**a**) *N. benthamiana* plants grew well in the experimental greenhouse field. The leaves of planted *N. benthamiana* plants (**b**, adaxial surface; **c**, abaxial surface. The pink arrows indicate dead whiteflies) attracted a significant number of whiteflies.

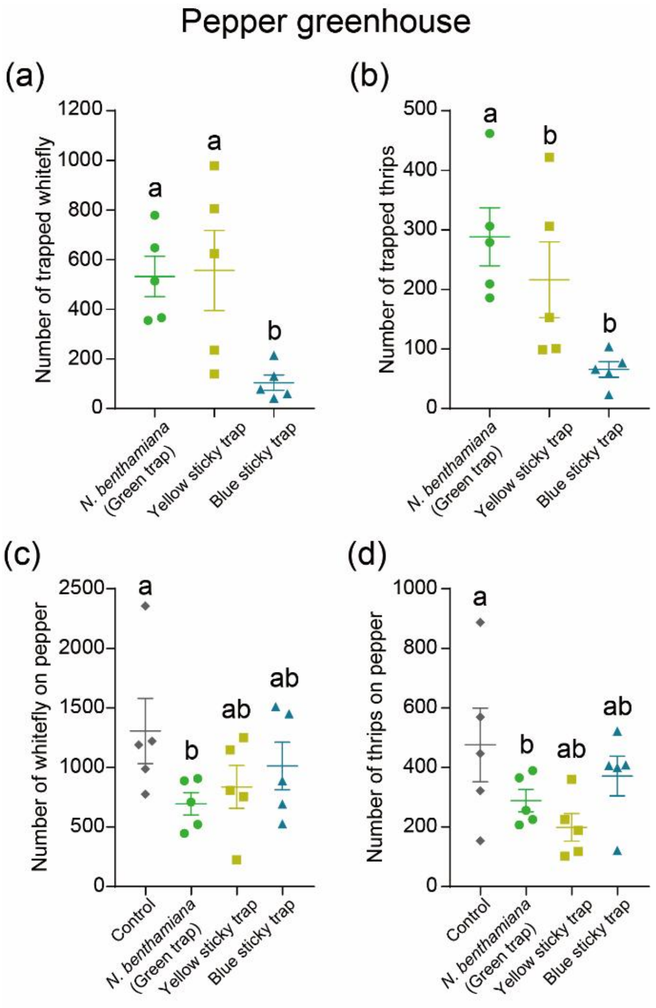


**Fig. S7.** ***N. benthamiana*** **plants as a substitute for commercial sticky traps in pepper greenhouse.** (**a**) The number of trapped whiteflies on the *N. benthamiana* plant, yellow sticky trap, and blue sticky trap in the pepper greenhouse. (**b**) The number of trapped thrips on the *N. benthamiana* plant, yellow sticky trap, and blue sticky trap in the pepper greenhouse. The number of whiteflies (**c**) and thrips (**d**) in pepper crops across the experimental plots are presented. Values are mean ± SEM, $n = 5$. One-way ANOVA followed by Fisher's least significant difference (LSD) test was used for significant difference analysis. Bars with different lowercase letters indicate significant differences between treatments at $p < 0.05$.

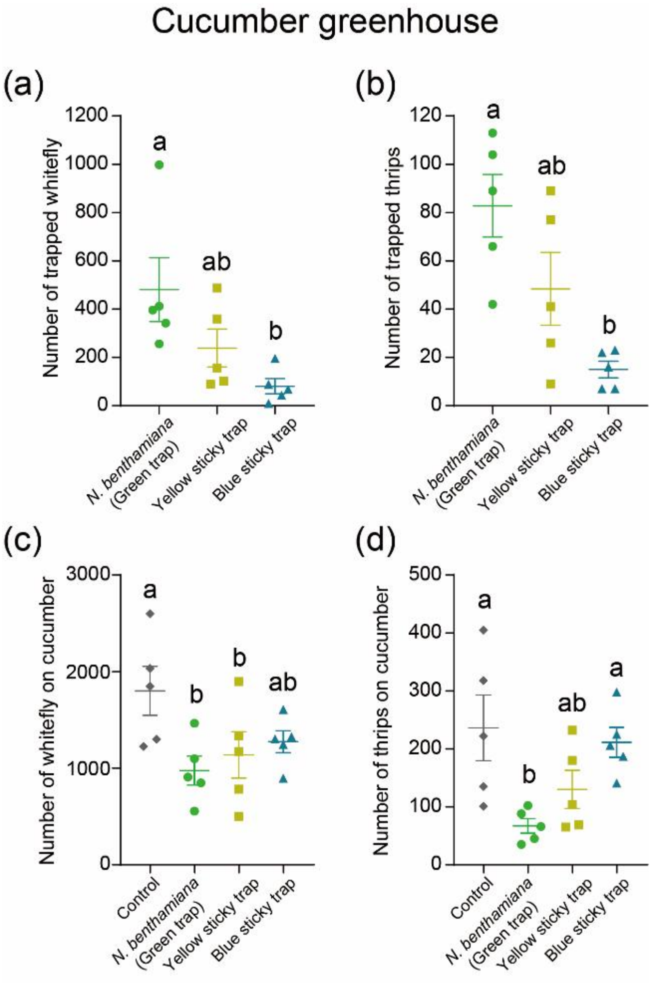


**Fig. S8. *N. benthamiana* plants as a substitute for commercial sticky traps in cucumber greenhouse.**

(**a**) The number of trapped whiteflies on the *N. benthamiana* plant, yellow sticky trap, and blue sticky trap in the cucumber greenhouse. (**b**) The number of trapped thrips on the *N. benthamiana* plant, yellow sticky trap, and blue sticky trap in the cucumber greenhouse. The number of whiteflies (**c**) and thrips (**d**) in cucumber crops across the experimental plots are presented. Values are mean ± SEM, $n$ = 5. One-way ANOVA followed by Fisher's least significant difference (LSD) test was used for significant difference analysis. Bars with different lowercase letters indicate significant differences between treatments at $p < 0.05$.

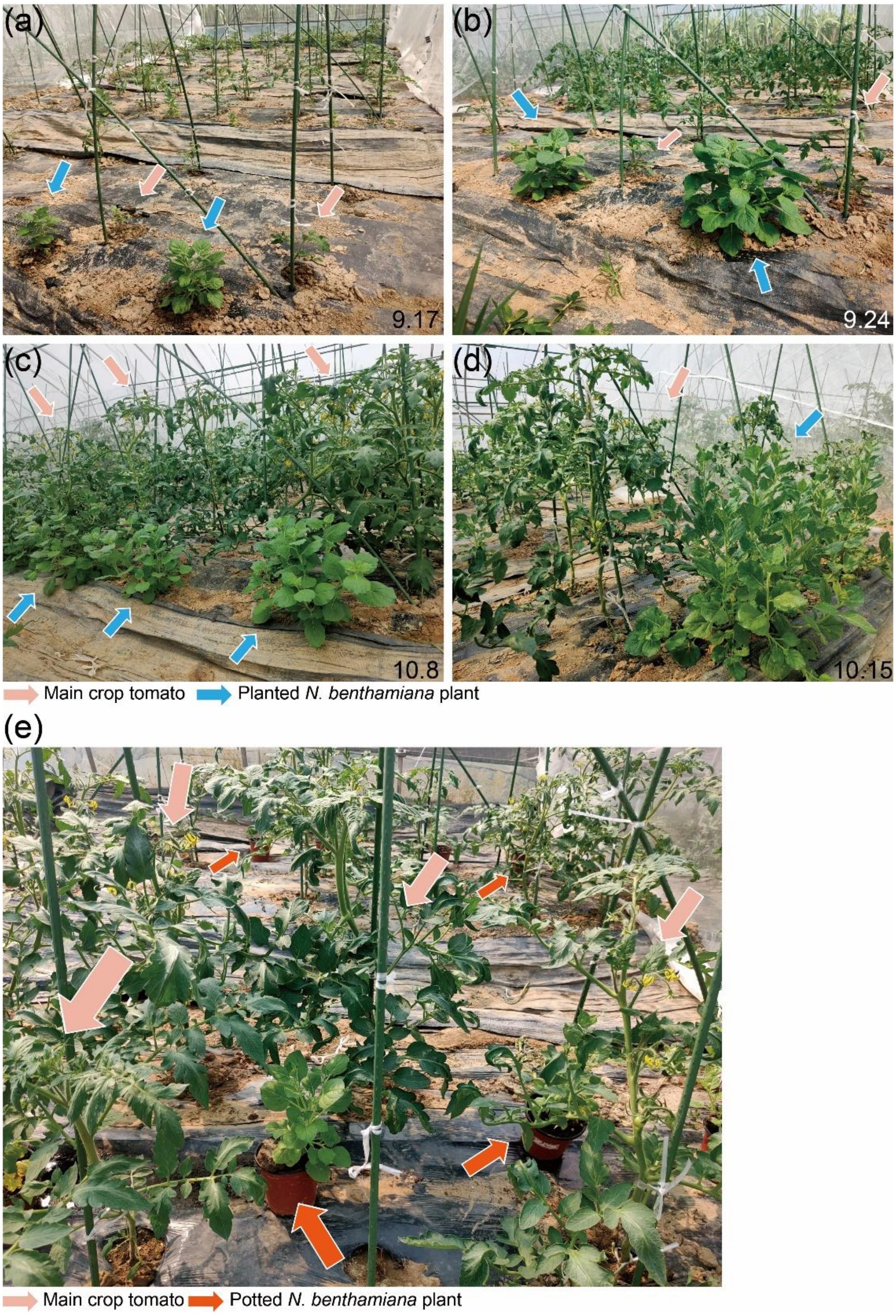


**Fig. S9. The status of field-grown *N. benthamiana* plants in the commercial greenhouse.**

The status of field-grown *N. benthamiana* plants and main crop tomatoes in the experimental greenhouse is present for different dates (**a** for 7$^{th}$ September; **b** for 24$^{th}$ September; **c** for 8$^{th}$ October; **d** for 15$^{th}$ October). The pink arrow indicates the main crop tomatoes and the blue arrow indicates the field-grown *N. benthamiana* plants. (**e**) The status of potted *N. benthamiana* plants in the commercial greenhouse. The pink arrow indicates the main crop of tomatoes. The orange arrow indicates the potted *N. benthamiana* plants.

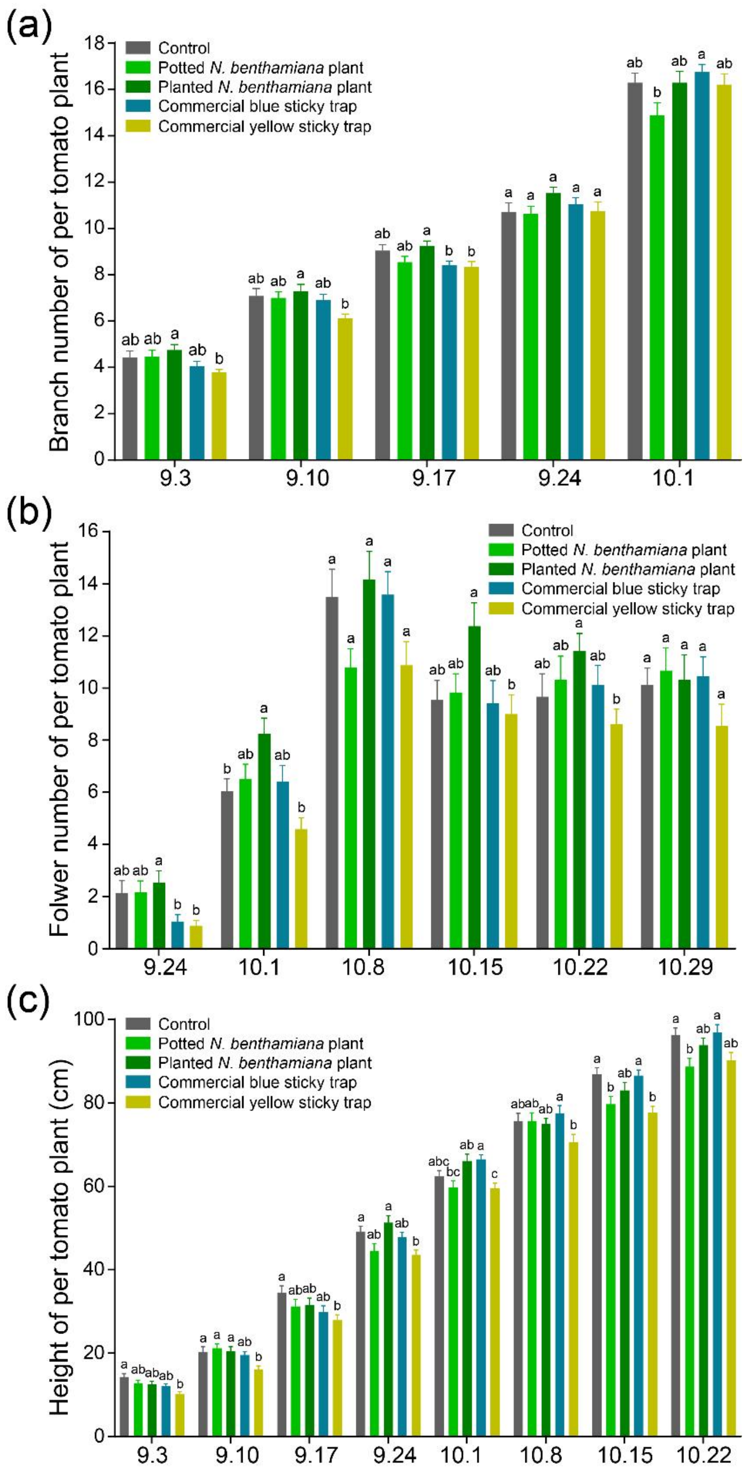


**Fig. S10. Traits of the main crop tomatoes in experimental plots with *N. benthamiana* plants and sticky traps.**

The branch number (**a**), plant height (**b**), and flower number (**c**) of tomatoes in different experimental plots are presented. Values are mean ± SEM, $n$ = 24. One-way ANOVA followed by Fisher's least significant difference (LSD) test was used for significant difference analysis. Bars with different lowercase letters indicate significant differences between treatments at $p < 0.05$.

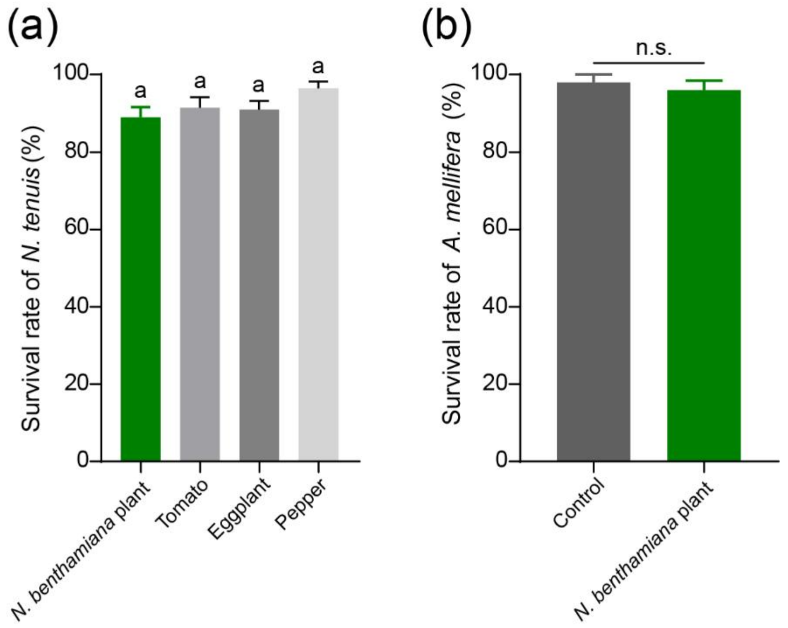


**Fig. S11. Effects of *N. benthamiana* plants on beneficial insects.**

(**a**) The survival rate of the natural enemy *N. tenuis* was evaluated on *N. benthamiana* plants, as well as tomatoes, eggplants, and peppers. (**b**) The survival rate of honeybees on *N. benthamiana* plants. Each *N. benthamiana* plant was enclosed within a nylon cage housing ten honeybees, and their survival was calculated. As a control, honeybees were placed within an empty cage. Values are mean ± SEM, $n = 10$ for **a**; $n = 5$ for **b**. One-way ANOVA followed by Fisher's least significant difference (LSD) test was used for significant difference analysis for **a**. Bars with different lowercase letters indicate significant differences between treatments at $p < 0.05$. Student's *t*-test (two-tailed) was used for significant difference analysis for **b**. n. s., not significant.
